# Developmental timing of programmed DNA elimination in *Paramecium tetraurelia* recapitulates germline transposon evolutionary dynamics

**DOI:** 10.1101/2022.05.18.492358

**Authors:** Coralie Zangarelli, Olivier Arnaiz, Mickaël Bourge, Kevin Gorrichon, Yan Jaszczyszyn, Nathalie Mathy, Loïc Escoriza, Mireille Bétermier, Vinciane Régnier

## Abstract

With its nuclear dualism, the ciliate *Paramecium* constitutes an original model to study how host genomes cope with transposable elements (TEs). *P. tetraurelia* harbors two germline micronuclei (MIC) and a polyploid somatic macronucleus (MAC) that develops from the MIC at each sexual cycle. Throughout evolution, the MIC genome has been continuously colonized by TEs and related sequences that are removed from the somatic genome during MAC development. Whereas TE elimination is generally imprecise, excision of ∼45,000 TE-derived Internal Eliminated Sequences (IESs) is precise, allowing for functional gene assembly. Programmed DNA elimination is concomitant with genome amplification. It is guided by non-coding RNAs and repressive chromatin marks. A subset of IESs is excised independently of this epigenetic control, raising the question of how IESs are targeted for elimination. To gain insight into the determinants of IES excision, we established the developmental timing of DNA elimination genome-wide by combining fluorescence-assisted nuclear sorting with next-generation sequencing. Essentially all IESs are excised within only one endoreplication round (32C to 64C), while TEs are eliminated at a later stage. We show that time, rather than replication, controls the progression of DNA elimination. We defined four IES classes according to excision timing. The earliest excised IESs tend to be independent of epigenetic factors, display strong sequence signals at their ends and originate from the most ancient integration events. We conclude that old IESs have been optimized during evolution for early and accurate excision, by acquiring stronger sequence determinants and escaping epigenetic control.

## Introduction

Transposable elements (TEs) have colonized the genomes of most living species and constitute a significant fraction of extant genomes, from a few percent in yeast (Bleykasten-Grosshans and Neuveglise 2011) to ∼85% in some plant genomes (Bennetzen and Park 2018). TEs are often considered as genomic parasites threatening host genome integrity, even though they can be a source of genetic innovation (Cosby et al. 2019; Capy 2021). Host defense pathways counteract the potentially detrimental effects of transposon invasion. In eukaryotes, small RNA (sRNA)-dependent post-transcriptional and transcriptional silencing mechanisms inactivate TE expression and transposition, both in germline and somatic cells (Ketting et al. 1999; Tabara et al. 1999; Zilberman et al. 2003; Brennecke et al. 2007). TE transcriptional inactivation is associated with heterochromatin formation, through DNA methylation and histone H3 methylation on lysine 9 (Deniz et al. 2019; Choi and Lee 2020). Another epigenetic mark, H3K27me3, also contributes to TE silencing in several species (reviewed in (Déléris et al. 2021)). Because of their remarkable germline-soma nuclear dualism (Prescott 1994; Cheng et al. 2020), ciliates are original unicellular eukaryotic models to study the dynamics of TEs within genomes, both at the developmental and evolutionary time-scales (Arnaiz et al. 2012; Hamilton et al. 2016; Kapusta et al. 2017; Sellis et al. 2021).

*Paramecium* species harbor one to four transcriptionally silent diploid germline micronuclei (MIC) (Görtz 1988) that coexist with a polyploid somatic macronucleus (MAC) responsible for gene expression. During sexual processes (conjugation of compatible reactive partners or a self-fertilization process called autogamy), the MICs undergo meiosis and transmit the germline genome to the diploid zygotic nucleus through fertilization and karyogamy (Bétermier and Duharcourt 2014). In the meantime the old MAC splits into ∼30 fragments that continue to ensure gene expression while new MICs and MACs differentiate from division products of the zygotic nucleus. The formation of a functional new MAC is essential to take over gene expression once old MAC fragments have disappeared from the cell. New MAC development covers two cell cycles after the zygotic nucleus is formed. During this period, massive genome amplification takes place within each developing MAC (also called anlagen) to reach the final endoduplication level of mature MACs (∼800C to 1600C in *P. tetraurelia*) (Preer 1976). Pioneering work from J. Berger suggested that at least 4 discontinuous peaks of DNA synthesis, corresponding to ∼5 doublings of DNA content, take place in anlagen until the first cell fission in *P. tetraurelia*, while 4.5 more doublings occur with a more continuous pattern during the second cell cycle (Berger 1973). Concomitantly with genome amplification, programmed genome rearrangements (PGR) remove ∼30% of germline DNA from the new MAC genome, going from ∼108-Mbp haploid genome size in the MIC to 72 - 75 Mbp in the mature MAC (Aury et al. 2006; Guérin et al. 2017; Sellis et al. 2021). Because eliminated DNA includes TEs and related sequences (Arnaiz et al. 2012; Guérin et al. 2017; Sellis et al. 2021), PGR in *Paramecium*, as in other ciliates, can be viewed as an extreme mechanism to inactivate TEs in the somatic genome.

Two types of germline sequences, referred to as “MIC-limited” DNA, are removed during PGR in *Paramecium* (Bétermier and Duharcourt 2014). At least 25% of the MIC genome, including DNA repeats (TEs, minisatellites), are eliminated imprecisely, alternatively leading to chromosome fragmentation (with *de novo* telomere addition to heterogeneous new MAC chromosome ends) or intrachromosomal deletions between variable boundaries (Baroin et al. 1987; Le Mouël et al. 2003; Guérin et al. 2017). In contrast, ∼45,000 Internal Eliminated Sequences (IESs) scattered throughout the germline genome (including inside coding sequences) are excised precisely, allowing for assembly of functional open reading frames (Arnaiz et al. 2012). *Paramecium* IESs are mostly short (93% <150 bp) non-coding sequences, with a damped sinusoidal size distribution extending from 26 bp to a few kbp. They are consistently flanked by two TA dinucleotides, one on each side, and leave a single TA on MAC chromosomes upon excision. Two independent studies, the first relying on the analysis of paralogous gene quartets originating from successive whole genome duplications in a single species, *P. tetraurelia* (Arnaiz et al. 2012), the other on phylogenetic analyses across 9 *Paramecium* species (Sellis et al. 2021), have made it possible to date ∼40% of *P. tetraurelia* IES insertions and define groups of old, intermediate and young IESs according to their evolutionary age. The oldest IESs, thought to have colonized the germline genome before divergence of *P. caudatum* and the *P. aurelia* clade, tend to be very short (26 to 30 bp) (Sellis et al. 2021). Several families of larger and younger IESs, some sharing homology with known *Paramecium* TEs, appear to have been mobile recently at the time-scale of *Paramecium* evolution: intermediate IESs were acquired after the divergence of *P. caudatum*, young IESs were gained after the burst of *P. aurelia* speciation. This is consistent with IESs being relics of ancestral TEs that have decayed during evolution through reduction in size and loss of coding capacity, while remaining under selection for precise excision from the MAC (Klobutcher and Herrick 1997; Dubois et al. 2012).

IES excision occurs through a “cut-and-repair” mechanism involving double-strand DNA cleavage around each flanking TA (Gratias and Bétermier 2003), followed by excision site closure through precise Non-Homologous End Joining (NHEJ) (Kapusta et al. 2011; Bétermier et al. 2014). Also depending on the NHEJ pathway, excised IESs were proposed to concatenate into end-to-end joined molecules before being eventually degraded (Allen et al. 2017), while those of sufficient length form transient covalently closed monomeric circles (Gratias and Bétermier 2001). Several components of the core IES excision machinery are known. The PiggyMac (Pgm) endonuclease, a catalytically active domesticated transposase (Baudry et al. 2009; Dubois et al. 2012) and its five PiggyMac-like partners, PgmL1 to PgmL5 (Bischerour et al. 2018), are essential for DSB introduction at IES ends. In the absence of Pgm, all IESs are retained in the anlagen and most imprecise DNA elimination is also impaired, except for ∼3 Mbp of germline sequences, the elimination of which seems to be Pgm-independent (Guérin et al. 2017). A specialized NHEJ factor, the Ku70/Ku80c (Ku) heterodimer also seems to be an essential component of the core endonuclease machinery: Ku is able to interact with Pgm, tethers it in the anlagen and licenses DNA cleavage at IES ends (Marmignon et al. 2014; Abello et al. 2020; Bétermier et al. 2020). *Paramecium* IES ends display a weak consensus (5’ <u>TA</u>YAGTNR 3’), too poorly conserved to serve as a specific recognition sequence for the endonuclease (Arnaiz et al. 2012). Additional epigenetic factors, including non-coding RNAs (ncRNAs) and histone modifications, control the recognition of eliminated DNA by the core machinery (Chalker et al. 2013; Bétermier and Duharcourt 2014; Allen and Nowacki 2020). According to the “scanning” model, sRNAs processed from meiotic MIC transcripts by Dicer-like proteins Dcl2 and Dcl3 are subtracted against old MAC sequences, resulting in the selection of a sub-population of scnRNAs covering the MIC-limited fraction of the germline genome (Lepère et al. 2008; Lepere et al. 2009). MIC-limited scnRNAs are thought to target elimination of their homologous sequences by pairing with TFIIS4-dependent non-coding nascent transcripts in the anlagen (Maliszewska-Olejniczak et al. 2015), thereby triggering H3K9 and K27 trimethylation by the histone methyltransferase Ezl1 (Lhuillier-Akakpo et al. 2014; Frapporti et al. 2019; Miró-Pina et al. 2022). The H3K9me3 and H3K27me3 heterochromatin marks are required for the elimination of TEs and ∼70% of IESs (Lhuillier-Akakpo et al. 2014; Guérin et al. 2017). A second population of sRNAs (called iesRNAs), produced from excised IES transcripts by the Dcl5 protein, was proposed to further assist IES excision (Sandoval et al. 2014; Allen et al. 2017). Both types of sRNAs appear to act synergistically. Indeed, while *DCL2/3* or *DCL5* knockdowns (KD) each impair excision of only a small fraction of IESs (∼7% in a *DCL2/3* KD, ∼5% in a *DCL5* KD) (Lhuillier-Akakpo et al. 2014; Sandoval et al. 2014) a triple *DCL2/3/5* KD inhibits excision of ∼50 % of IESs coinciding with the set of TFIIS4-dependent IESs (Swart et al. 2017), which we refer to as ncRNA-dependent IESs.

While our knowledge of the molecular mechanisms involved in the epigenetic control and catalysis of PGR in *P. tetraurelia* has increased over the past decade, little is known about the relative timing of DNA replication and PGR during MAC development. Molecular data obtained for a handful of IESs have suggested that excision starts after several endoreplication rounds have already taken place in the anlagen (Bétermier et al. 2000), but that not all IESs are excised at exactly the same time (Gratias and Bétermier 2001). Here, we have investigated at the genome-wide level the elimination timing of all 45,000 IESs and other MIC-limited sequences, including TEs. We set up a fluorescence-assisted nuclear sorting (FANS) procedure to selectively purify anlagen at different ploidy levels during autogamy and sequence their DNA. We show that essentially all IESs are excised within one endoreplication round (between 32C and 64C) and that most TE elimination and formation of new MAC chromosome ends take place following one additional replication round. We define four classes of “very early”, “early”, “intermediate” and “late” IESs according to their excision timing. We provide evidence that very early excised IESs tend to be short, amongst the oldest in the genome, and independent of sRNAs and heterochromatin marks for excision. We also found that they display stronger nucleotide signals at their ends. Our data strongly support an evolutionary scenario in which *Paramecium* IESs have evolved from germline TEs under mechanistic constraint to be excised early and accurately from the somatic genome, independently of ncRNAs or heterochromatin marks.

## Results

### A Fluorescence-Activated Nuclear Sorting (FANS) strategy to purify new MACs

To selectively purify developing new MACs during an autogamy time-course (tc), we adapted a published flow cytometry procedure that was first set up to sort anlagen from old MAC fragments at a late developmental stage, when the two types of nuclei can clearly be distinguished based on their size and DNA content (Guérin et al. 2017). Because at early stages (DEV1 and DEV2, see Methods), anlagen and old MAC fragments have similar sizes (Fig. 1A), we selectively labeled the new MACs using a specific α-PgmL1 antibody raised against a component of the IES excision machinery (Bischerour et al. 2018). We first confirmed that immunofluorescence staining of whole cells yielded a strong and specific signal in the anlagen throughout DEV1 to DEV4 (Fig. 1A; Supplemental Fig. S1A,B,C), which corresponds to the time-window (T3 to T30) when programmed DSBs are detected at IES boundaries (Gratias et al. 2008; Baudry et al. 2009). Using the α-PgmL1 antibody to label unfixed nuclei harvested at DEV1 and DEV2 during an autogamy time-course of wild-type (wt) cells, we confirmed that PgmL1 labeling can be used to separate anlagen from old MAC fragments using flow cytometry (Fig. 1B and Supplemental Fig. S1D,E).

**Figure 1.**
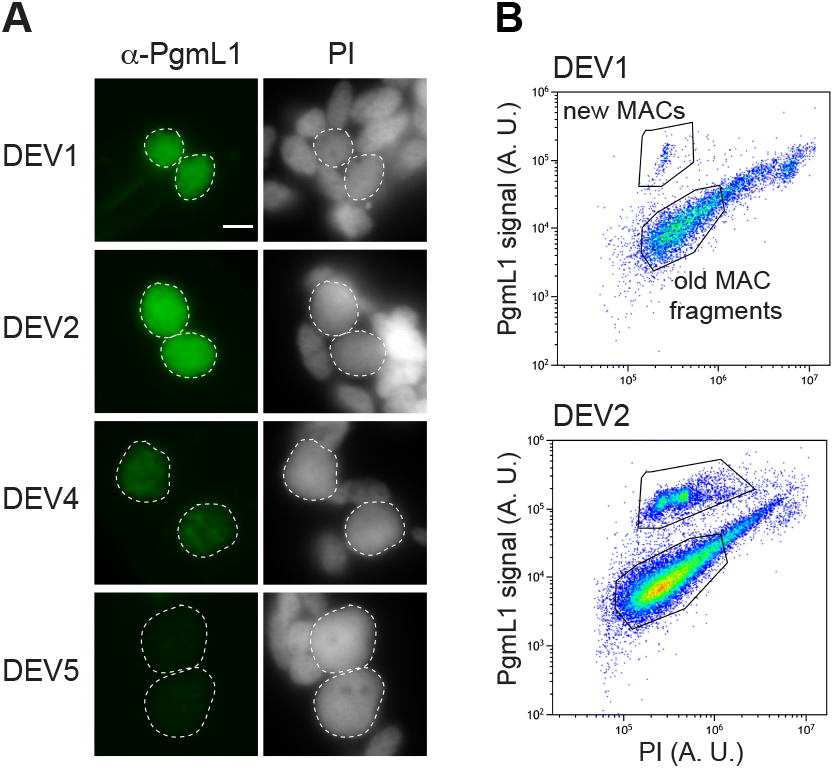
PgmL1 immunostaining during autogamy. (*A*) Whole cell immunostaining at different stages of autogamy time-course 1 (tc1). New MACs and fragments are counterstained with Propidium Iodide (PI). Developing MACs are surrounded by a white dotted line. Scale bar is 5 µm. Developmental stages (DEV1 to 5) are defined in Methods. (B) Flow cytometry analysis of immunostained nuclei at the DEV1 and DEV2 stages of autogamy time-course 2 (tc2). Following gating of total nuclei (see Supplemental Fig. S1D), the population of new MACs was separated based on their PgmL1 signal. The PI axis is indicative of DNA content. A. U.: arbitrary units in log scale.

### Most IES excision takes place within one round of replication

We used FANS to purify new MACs at DEV1 to DEV4 (Fig. 2A). The distribution of Propidium Iodide (PI) fluorescence intensities revealed a series of three discrete peaks from DEV1 to DEV3, indicative of the presence of nuclear populations with a defined DNA content. We calculated the C-value for each peak (see Methods and Supplemental Figure S2) and further evaluated their C-level using an estimated 1C-value of ∼100Mb for the unrearranged *P. tetraurelia* MIC genome (Guérin et al. 2017; Sellis et al. 2021). We found C-levels of ∼32C, ∼64C and ∼128C (tc4 in Supplemental Table S1), which is consistent with successive whole genome replication rounds. The ∼32C peak at DEV1 indicates that 4 replication rounds have been completed at this stage. Two more rounds take place until DEV3. At DEV4, which is the final stage where PgmL1 staining can be detected, we observed an enlargement of the ∼128C peak, indicative of a mixed population with a more variable amount of DNA. This could reflect a switch to an asynchronous replication mode, as previously reported (Berger 1973).

**Figure 2.**
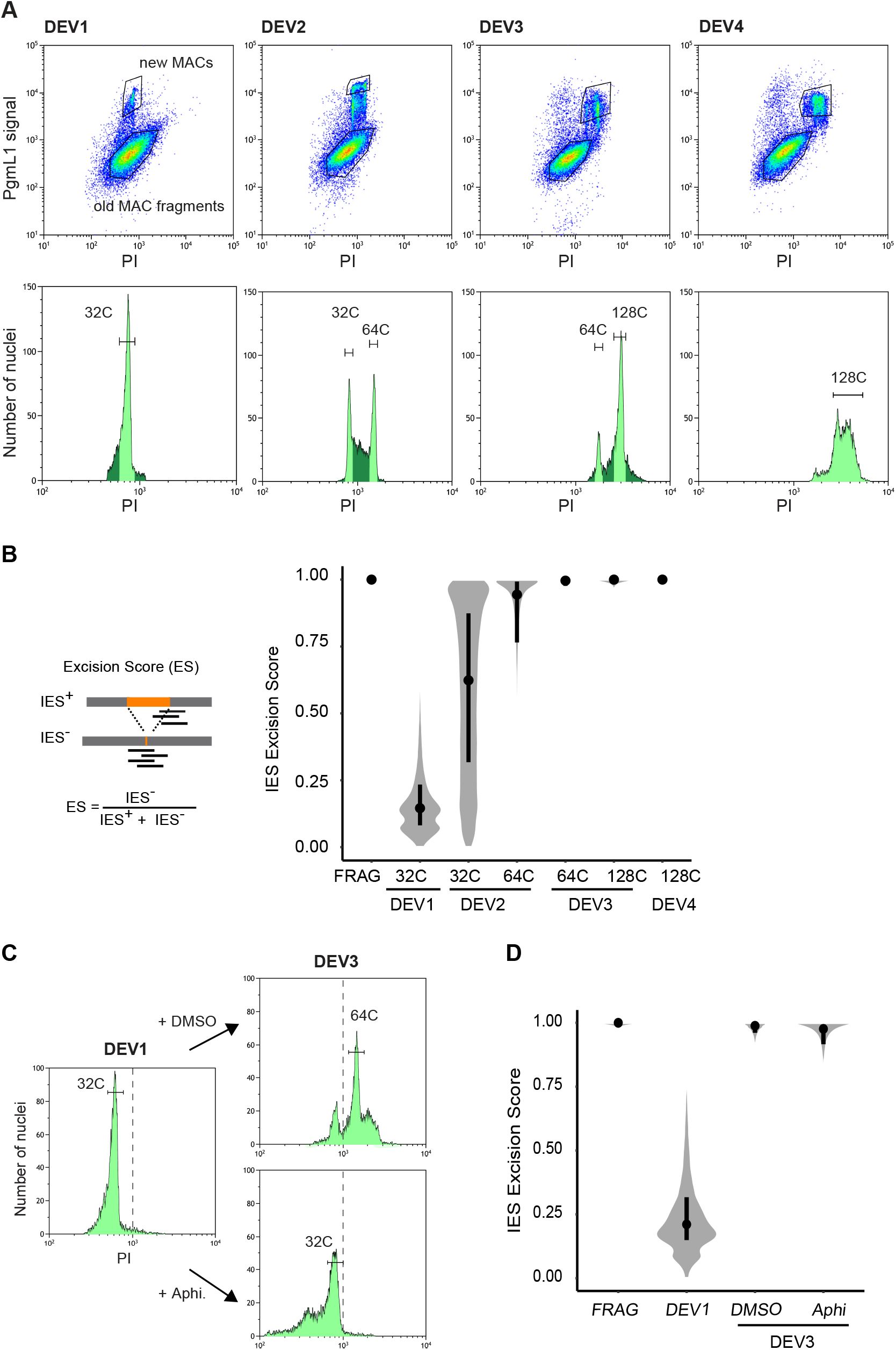
IES excision kinetics and endoreplication. (*A*) Flow cytometry sorting of nuclei during the different stages of an autogamy time-course (tc4). Upper panels: Plots of PgmL1 fluorescence intensity (y-axis) versus PI fluorescence intensity (x-axis) for nuclei collected at different developmental stages. Lower panels: Histograms of PI-stained nuclei gated in the upper panel. Sorted new MAC peaks are indicated by light green shading. The C-level for each sorted peak was estimated as described in Supplemental Table S1 and is indicated above each peak. For DEV4 nuclei, the whole PgmL1-labeled population was sorted (light green), but the major peak was used for calculation of the C-level. As a control, old MAC fragments were sorted from the DEV1 stage. (*B*) Distribution of IES Excision Scores (ES) in the different sorted new MAC populations. Samples are named according to the developmental stage (DEV1 to DEV4 from tc4) and the C-level of the sorted population. A schematic representation of the IES+ and IES-Illumina sequencing reads that were counted to calculate the ES is presented on the left. An ES of 0 or 1 corresponds to no or complete IES excision, respectively. The black dot is the median and the vertical black line delimitates the second and third quartiles. (*C*) Flow cytometry sorting of nuclei following aphidicolin treatment. PI histograms of PgmL1-labelled nuclei are presented for each stage or condition (DEV1, DEV3 DMSO and DEV3 Aphi). The C-level for the indicated peaks was estimated as described in Supplemental Table S1. For each stage, all PgmL1-labelled nuclei were sorted. Old MAC fragments were sorted as a control from the DEV3 DMSO nuclear preparation. The dotted line is indicative of a PI value of 103. (*D*) ES distribution in the sorted new MAC populations in the aphidicolin time-course. Sample names correspond to the sorted samples shown in *C*.

We further sorted the populations of nuclei issued from each peak (Fig. 2A) and extracted their DNA for deep sequencing (tc4 in Supplemental Table S2). Thanks to the absence of old MAC contamination (See Methods and Supplemental Fig. S3), molecules lacking an inserted IES (designated IES^−^) correspond to *de novo* excision junctions. Therefore, this procedure makes it possible for the first time to calculate a real excision score (ES) for each of the 45,000 IESs (Fig. 2B). At DEV1 ∼32C, only few IESs have been excised, with a median ES value of 0.15. The median ES rises to 1 at DEV3 ∼64C, indicating that nearly all IESs are excised within one round of replication. To investigate whether the 5^th^ endoreplication round itself is mandatory for DNA elimination, we treated autogamous cells with aphidicolin, a specific inhibitor of eukaryotic DNA polymerase α, after they reached DEV1 ∼32C (Fig. *2C*). Comparison of the flow cytometry profiles confirmed that the new MACs of control cells (+DMSO) have undergone their 5^th^ replication round at DEV3, while those of aphidicolin-treated cells are blocked at ∼32C. We further sorted anlagen from the DEV1, DEV3 DMSO and DEV3 Aphi samples for DNA sequencing (Supplemental Table S2). As shown in Fig. 2D, the median ES for untreated new MACs at DEV3 is 0.99, confirming the completion of IES excision. For aphidicolin-treated anlagen, the median ES is 0.98, indicating that inhibiting the 5^th^ endoreplication round does not impair IES excision.

### Imprecise elimination is delayed relative to IES excision

To strengthen our analysis of the timing of DNA elimination, we included sorted samples from 4 replicate time-course experiments (Supplemental Fig. S4A,B; Supplemental Tables S1,S2). The resulting ES distributions strengthen our conclusion that IES excision takes place between DEV1 ∼32C and DEV3 ∼64C (Fig. 3A). We used the same sequencing data (Supplemental Table S2) to study the timing of imprecise DNA elimination during MAC development. Because this process yields heterogeneous MAC junctions, preventing us from calculating an ES, we analyzed sequencing data by read coverage. We found that IES coverage dropped between DEV1 ∼32C and DEV3 ∼64C (Fig. 3B), consistent with the excision profile obtained by ES calculation (Fig. 3A). The TE sequencing coverage exhibited a delayed decrease, with a drop starting at DEV3 ∼64C, also observed for the whole MIC genome coverage. Because sequence coverage analysis cannot discriminate between intra- and extra-chromosomal molecules, we examined whether imprecise elimination could have taken place before the drop in coverage. Imprecise elimination of MIC-specific sequences (containing TEs) is accompanied by chromosome fragmentation and formation of *de novo* telomeric ends (Le Mouël et al. 2003). We observed that telomeric reads only increase at DEV3 ∼64C (Fig. 3C), supporting the idea that imprecise elimination does not begin before DEV3. Of note, the whole MIC genome coverage at DEV4 ∼128C is still higher than the genome coverage of fragments (which harbor a fully rearranged genome), indicating that imprecise elimination is not totally completed in the new MAC at this stage.

**Figure 3.**
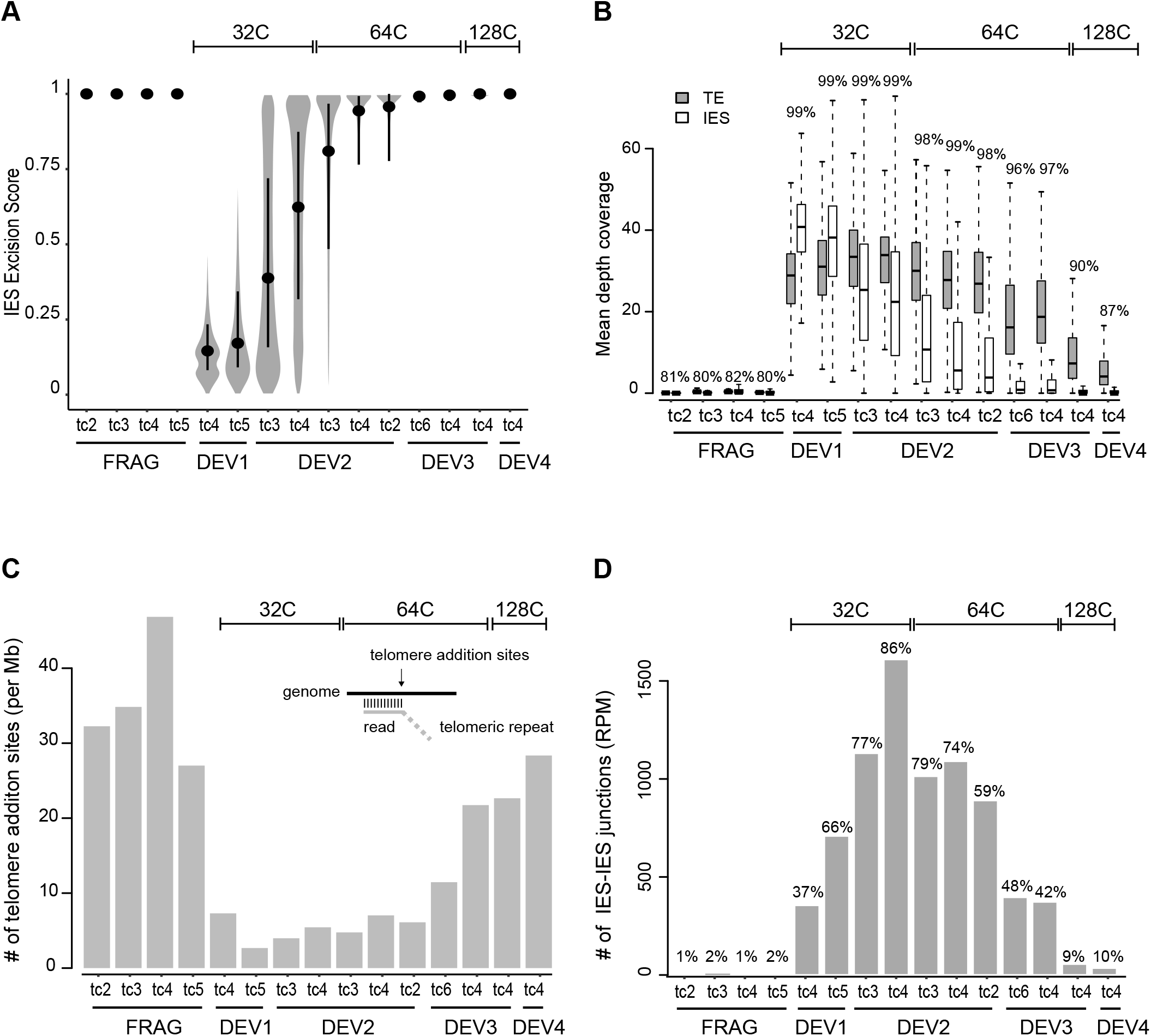
Kinetics of precise IES excision and imprecise DNA elimination. (*A*) Distribution of ESs in all samples. Samples are named and ordered according to developmental stage (DEV1 to DEV4), time-course (tc2, tc3, tc4, tc5, tc6) and C-level (indicated above the plot). For each time-course, old MAC fragments (FRAG) were also sorted as controls. The black dot is the median and the vertical black line spans the second and third quartiles. (*B*) TE and IES coverage during autogamy. The mean depth coverage distribution is represented as a boxplot. For each dataset described in *A*, the grey boxplot shows TE coverage, and the white, IES coverage. The percentage of the MIC genome covered by the sequencing reads is indicated above each pair of boxplots. (*C*) Abundance of telomere addition sites during autogamy. The scheme above the bars illustrates the method for detection of telomere addition sites using the sequencing data. For each dataset, the bar shows the normalized number (per Mb of reads mapped) of detected telomere addition sites. (*D*) Quantification of IES-IES junctions. All putative molecules resulting from ligation of excised IES ends (See Supplemental Fig. S5A,B) are counted and normalized using sequencing depth.The percentage of IESs involved in at least one IES-IES junction is indicated above the barplot.

### Genome wide detection of transient IES-IES junctions

The purity of the FANS-sorted anlagen increases our ability to detect transient molecules produced during IES excision. We developed a new bioinformatic method to quantify IES-IES junctions (Fig. 3D) and confirmed that new MACs indeed contain molecules corresponding to the two expected types of excised junctions (single-IES circles and multi-IES concatemers) (Supplemental Fig. S5A). Because the normalized count of IES-IES junctions is maximal at DEV2 ∼32C (Fig. *3D*), the stage at which ESs dramatically increase (Fig. *3A*), we conclude that IES-IES junctions are formed concomitantly with MAC junctions, but are still detected at ∼64C DEV3, when IES excision is completed. This confirms, at the genome-wide level, that excised IES products are not degraded immediately and persist in the new MACs (Bétermier et al. 2000). Our data also reveal that the vast majority of IESs (97.2% considering all datasets and 86% at DEV2 ∼32C) are involved in the formation of IES-IES junctions (Fig. 3D). Based on read counts, single-IES circles represent fewer than 2% of excised IES junctions, indicating that concatemers are the major products of IES-end joining following excision (Supplemental Fig. S5B). This may be attributed to the size distribution of IESs, 93% of which are shorter than 150 bp (Arnaiz et al. 2012), the persistence length of dsDNA, and would therefore inefficiently form monomeric circles. Consistently, the size distribution of single-IES circles is centered around 200 bp (Supplemental Fig. S5C), indicating that only the longest IESs self-circularize.

### Sequential timing of excision is associated with specific IES features

We further used the excision score for the ∼45,000 IESs across all samples to group IESs into 4 clusters, according to their excision timing (“very early”, “early”, “intermediate”, “late”; Fig. 4A; Supplemental Fig. S6A; Supplemental Table S3). Very early IESs are almost all excised at DEV2 ∼32C, while excision of most late IESs takes place between DEV2 ∼64C and DEV3 ∼64C. Detection of IES-IES junctions follows the same excision timing: IESs from the very early and early clusters contribute to the majority of junctions detected at the earliest developmental stages, while IESs from the intermediate and late clusters become dominant at late developmental stages (Supplemental Fig. S6B). It has been previously observed that the excision machinery sometimes generates different types of errors (Supplemental Fig. S7A) (Duret et al. 2008; Bischerour et al. 2018). At the stage where all IESs are completely excised (DEV4 ∼128C), we observe 7-fold fewer excision errors for very early relative to late excised IESs (Fig. 4B), indicating that very early IESs are much less error-prone. Of note, the maximum of excision errors during the excision time-course never exceeds the error level observed in old MAC fragments (Supplemental Fig. S7B,C).

**Figure 4.**
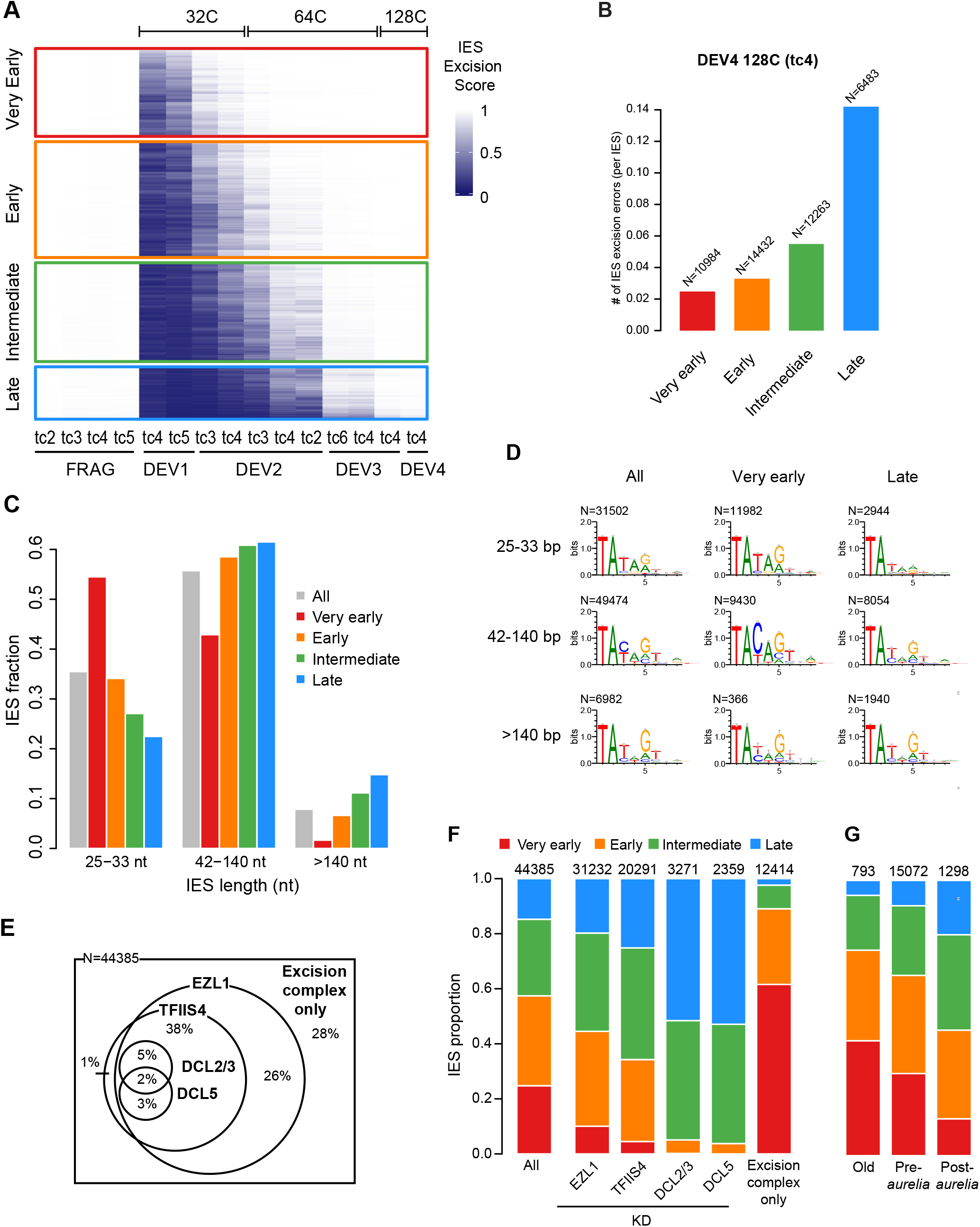
Excision timing defines IES classes with different characteristics. (*A*) Heatmap of ESs for all IESs. IESs are sorted by hierarchical clustering, each row corresponding to one IES. The ES is encoded from 0 (dark blue, no excision) to 1 (white, complete excision). IESs are separated in 4 classes according to their excision profile by K-means clustering of their ES (very early: N=10,994, early: N=14,490, intermediate: N=12,353, late: N=6,548). (*B*) Abundance of IES excision errors in the four excision profile groups counted in the DEV4 128C (tc4) sample. Only excision error types that are theoretically unbiased by IES length are considered (external, overlap and partial external, see Supplemental Fig. S7*A* for the definition of IES excision errors). The number of IESs in each excision profile group is indicated above the bars. (*C*) IES fraction for IES length categories in the four excision profile groups compared to all IESs. (*D*) Sequence logos of the 8 bases at IES ends for all IESs and IESs belonging to the very early and late clusters. IESs are grouped in three length categories as described in *C*. (*E*) Venn diagram showing how the 44,385 reference IESs are distributed according to their dependency on Ezl1, TFIIS4, Dcl2/3 and Dcl5 for excision, using an RNAi approach. The group “excision complex only” represents IESs that do not depend on any of these factors but do depend upon Pgm. The Venn diagram has been simplified to display only overlaps representing more than 1% of the total number of IESs. (*F*) IES proportions in the 4 groups of excision profiles for the datasets defined in *E*. The numbers above the barplots indicate the number of IESs in each dataset. “All” is the random expectation for all IESs. (*G*) IES proportions in the 4 groups of excision profiles relative to the age of IES insertion during evolution of the *Paramecium* lineage. Old: insertion predating the divergence between *P. caudatum* and the *P. aurelia* clade. Pre-*aurelia*: insertion before the radiation of the *P. aurelia* complex. Post-*aurelia*: insertion after the radiation of the *P. aurelia* complex (Sellis et al. 2021).

With regard to genomic location, we found that late IESs are under-represented in genes, particularly in coding sequences (CDS) versus introns while the inverse trend is observed for very early and early IESs (Supplemental Fig. S8A). We also observed a strong enrichment of late excised IESs and a depletion of very early and early IESs at the extremities of MAC scaffolds, which is consistent with these regions being gene-poor (Supplemental Fig. S8B). Under-representation of late excised IESs within genes might be explained by a selective pressure for accurate excision to avoid the formation of non-functional ORFs.

As for IES intrinsic properties, we detected an impressive size bias between IESs from the different clusters, with the very early excised IESs tending to be much shorter than expected from the global IES size distribution. In contrast, short IESs are under-represented among late IESs (Fig. 4C; Supplemental Fig. S9). We then examined whether IESs have different sequence properties at their ends depending on the cluster they belong to. Because the consensus of IES ends varies at positions 3, 4 and 5 (position 1 being the T from the TA boundary) as a function of IES length (Swart et al. 2014), we compared sequence logos of IES ends in the different clusters according to IES size categories (Fig. 4D and Supplemental Fig. S10A,B). For 25 to 33-bp IESs, we found an over-representation of the TATAG boundary among very early IESs compared to late IESs. The increase of G frequency at the 5^th^ base position is particularly striking (62% vs 35% for very early compared to late IESs). For 42 to 140-bp IESs, we observed an even stronger sequence bias with an over-representation of the TACAG boundary among very early IESs, the increase of the C frequency at the third position being strongly significant (77% *vs* 30% for very early *vs* late IESs, respectively). We conclude that very early IESs shorter than 140 bp tend to exhibit a stronger nucleotide sequence signal at their ends than late excised IESs. No significant sequence difference between very early and late IESs was observed for longer IESs (>140 bp).

We further studied the link between excision timing and dependence upon known factors involved in the epigenetic control of IES excision (Fig. 4E,F; Supplemental Fig. S11). We found an under-representation of the very early excised cluster amongst the subset of IESs, whose excision depends on the deposition of H3K9me3 and H3K27me3 marks (*i*.*e*. IESs retained in *EZL1* KD) (Lhuillier-Akakpo et al. 2014). A similar bias was observed among IESs depending on the production of ncRNA from the anlagen (retained in *TFIIS4* KD) (Maliszewska-Olejniczak et al. 2015) and was exacerbated for sRNA-dependent IESs (retained in *DCL2/3* or *DCL5* KDs) (Sandoval et al. 2014; Lhuillier-Akakpo et al. 2014). In the *DCL* datasets, IESs from the very early cluster are totally absent, while IESs from the intermediate and, even more strikingly, the late clusters are over-represented. In contrast, very early excised IESs are strongly enriched (∼60%) among the 12414 IESs that are excised independently of the above factors (“excision complex only”). Considering the overlap between IES dependencies (Fig. 4E), our data indicate that IESs depending on known heterochromatin-targeting factors tend to take longer to be excised during MAC development.

Finally, we examined the relationship between IES evolutionary age (Sellis et al. 2021) and excision timing (Fig. 4G). Our data indicate that old IESs that invaded the *Paramecium* genome before the divergence of the *P. caudatum* and *P. aurelia* lineages tend to be precociously excised. Reciprocally, we observed that the younger the IESs, the later their excision during MAC development.

### Identification of new IESs in MIC-limited regions

The presence of IESs in MIC-limited regions was reported previously but only a few examples have been described (Le Mouël et al. 2003; Duharcourt et al. 1998). We took advantage of the sequencing data we obtained during the course of MAC development to pinpoint precise excision events within late eliminated regions, therefore identifying new *bona fide* IESs (see Supplemental Methods). Their excision coud be transiently observed before their surrounding DNA is completely eliminated. We could identify a set of 167 IESs localized in imprecisely eliminated regions (Imp IESs) and 226 IESs located inside IESs (Internal IESs) from the reference set (Supplemental Fig. S12A,B; Supplemental Tables S4, S5). We found that Imp IESs are strongly biased towards short sequences while Internal IESs present no major difference in size compared to the reference IESs (Supplemental Fig. S12C). Concerning the dependency on heterochromatin marks, we found two contrasting situations for internal IESs: 26% are completely excised in Ezl1-depleted relative to control cells (IES Retention Score (IRS) ∼0) and 20% are strongly retained (IRS ∼1) (Supplemental Fig. S12D; Supplemental Table S6). The IRS is strongly correlated with IES size, Ezl1-independent IESs being much shorter than the dependent ones, a bias already observed for the reference IES set (Lhuillier-Akakpo et al. 2014). For their related encompassing IESs (n=223), we observed that 94% are significantly retained in Ezl1-depleted cells, consistent with their late excision timing. Interestingly, we found that most Imp IESs are independent of Ezl1-mediated heterochromatin marks for their excision (Supplemental Fig. S12D). Moreover, we noticed that the most independent are the shortest, with a breakpoint size of 33 nt (Supplemental Fig. S12E).

Taken together, our data confirm that IESs are scattered all along MIC chromosomes including MIC-limited regions, as previously hypothesized (Sellis et al. 2021). Internal IESs exhibit similar characteristics to those of the reference IES set in terms of length and epigenetic control, and therefore might share the same evolutionary history. The properties of Imp IESs, however, raise the question of their origin. Most of IESs are derived from TEs, but a previous report showed that genomic fragments can be co-opted to become IESs (Singh et al. 2014). We could therefore speculate that Imp IESs are excision-prone genomic fragments recognized by the excision machinery independently of histone mark deposition.

## Discussion

### Developmental timing of sequential DNA elimination

The present study aimed at unravelling the links between two intertwined DNA-driven mechanisms underlying somatic nuclear differentiation in *Paramecium:* genome amplification and programmed DNA elimination. Setting up the FANS procedure allowed us to demonstrate that genome amplification during MAC development is an endocycling process, defined as alternating S and G phases without mitosis (Lilly and Duronio 2005). In the time-window during which PGR take place, we identified three peaks representing discrete new MAC populations differing in their DNA content. The estimated C-levels of the first two peaks (∼32C, ∼64C) are consistent with them resulting from successive whole-genome doublings, based on an approximate average 1C-value of ∼100 Mbp (Supplemental Table S1). The range of C-levels obtained for the third peak fits less well with ∼128C, which can be explained by ongoing massive DNA elimination between ∼64C and ∼128C causing variability in the actual 1C-value of the anlagen.

We determined at an unprecedented resolution the timing of IES excision and imprecise elimination genome-wide, across successive endoreplication cycles (Fig. 5A). Our data show that DNA elimination is an ordered process. We found that most IESs are excised between DEV1 ∼32C and DEV3 ∼64C under standard conditions, while imprecise elimination only starts at DEV3 ∼64C. We established four classes of IESs according to their excision timing (very early IESs are eliminated at DEV2 ∼32C, early and intermediate IESs between DEV2 ∼32C and DEV2 ∼64C, while late IESs are excised at DEV3 ∼64C). By arresting cells at ∼32C with aphidicolin, we also demonstrated that time, rather than the process of replication itself, controls the progression of IES elimination once it has started. Our data thus indicate that the excision machinery is recruited to its chromatin target independently of replication fork passage. As previously suggested (Bétermier et al. 2000), the 3 to 4 endoreplication rounds preceding IES excision might still contribute to remodel chromatin and make it a suitable substrate for the excision machinery. In support of this hypothesis, several chromatin remodelers and histone chaperones are known to control PGR in *P. tetraurelia* (Ignarski et al. 2014; de Vanssay et al. 2020; Singh et al. 2022), but the temporal and mechanistic details of their action remain to be precisely understood.

**Figure 5.**
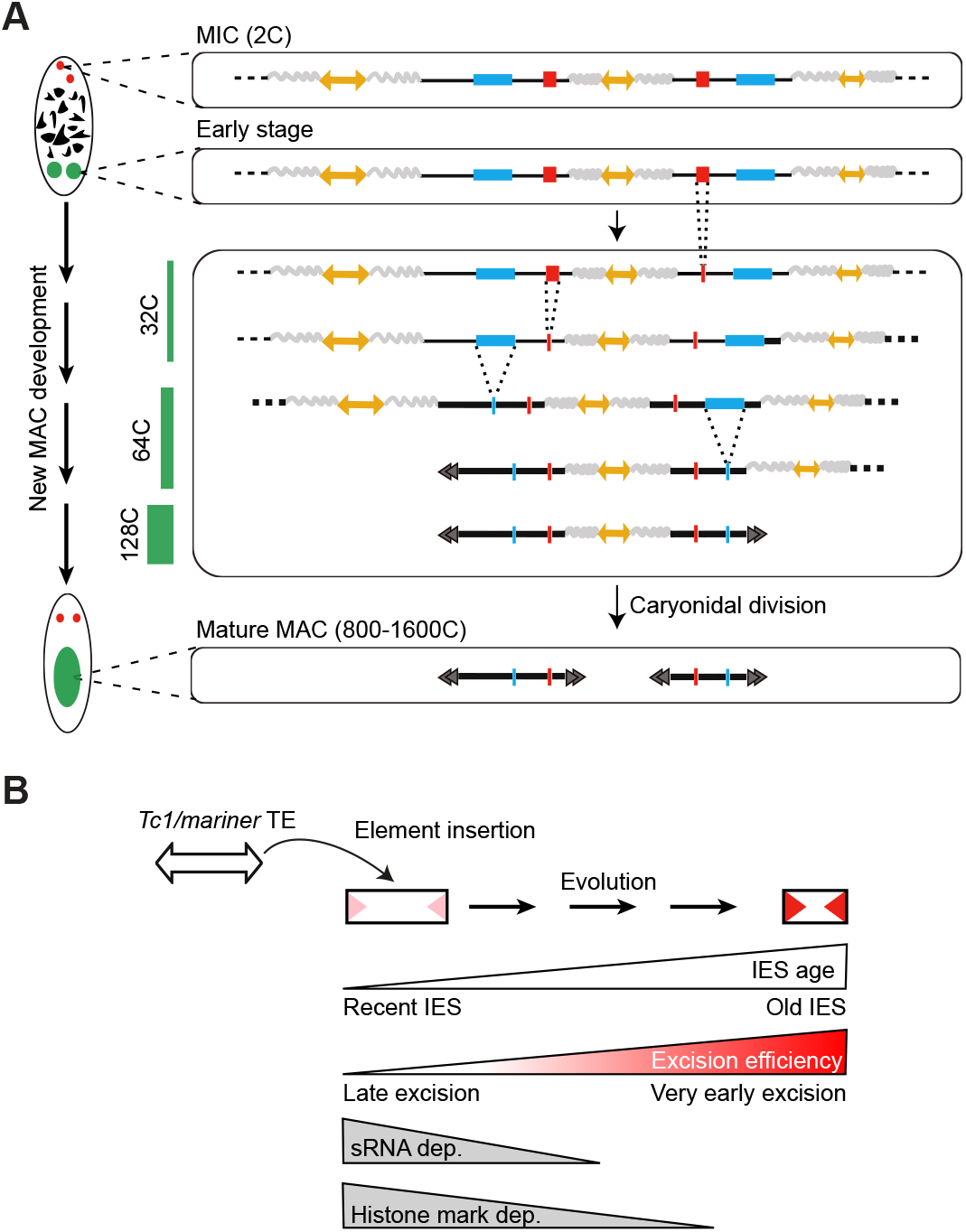
Schematic view of DNA elimination timing in *Paramecium* and model for IES evolution. (*A*) Schematic representation of the relative timing of DNA amplification and PGR during new MAC development. Double-headed yellow arrows stand for TEs and the surrounding wavy lines for their flanking imprecisely eliminated sequences. Red and blue boxes represent very early and late excised IESs, respectively. The endoreplication level (C-level) is indicated as a green bar on the left. The black double arrowheads schematize the telomeric ends of MAC chromosomes. At each step of PGR, only one representative copy of the new MAC genome is drawn. **(*B*)** Model for evolutionary optimization of IESs. Old IESs have become independent of sRNAs and histone mark deposition for their excision. They have acquired strong sequence information at their ends (red arrowheads), promoting their efficient excision.

Such a sequential DNA elimination program may provide *Paramecium* with an original mechanism to fine-tune zygotic gene expression during MAC development. IESs could block zygotic gene expression as long as they are retained within coding sequences or in gene regulatory regions, as suggested for the *PTIWI10* gene (Furrer et al. 2017). An even more sophisticated regulatory scheme could be proposed with a first IES excision event turning on an IES-containing anlagen-specific gene that would later be turned off by a second IES excision event. Another type of regulation was proposed previously (Sellis et al. 2021) for genes located inside IESs or embedded in imprecisely eliminated MIC-limited regions: such germline-specific genes may be expressed from the new MAC until their encompassing DNA is removed from the genome. In the future, monitoring the timing of PGR in other *Paramecium* species and annotating the sequential versions of the rearranged genome should allow us to assess whether temporal control of IES excision has been conserved during evolution and to what extent it may contribute to gene regulation.

### DNA elimination timing reveals evolutionary optimization of TE-derived sequences for efficient excision

Diverse TE families, including DNA and RNA transposons, have colonized the *Paramecium* germline genome and are mostly eliminated in a imprecise maner from the new MAC during PGR (Arnaiz et al. 2012; Guérin et al. 2017). We showed that imprecise elimination of TEs and other MIC-limited regions occurs at a late stage during MAC development (DEV3 to DEV4). TE elimination was previously shown to depend upon scnRNA-driven deposition of H3K9me3 and H3K27me3 histone marks, which are enriched on TEs at a developmental stage corresponding to DEV2 and accumulate in the anlagen up to DEV3 (Lhuillier-Akakpo et al. 2014; Frapporti et al. 2019). The deposition of heterochromatin marks thus appear to be a late process, which might explain why TEs and other MIC-limited sequences are eliminated at a late developmental stage.

*Paramecium* IESs have been proposed to originate from *Tc1/mariner* TEs, a particular family of transposons that duplicate their TA target site upon integration into the germline genome. Target site duplication generates potential Pgm cleavage sites at TE boundaries that could allow integrated TEs to be excised precisely from the developing MAC and thus become IESs (Arnaiz et al. 2012; Sellis et al. 2021). During evolution, IESs have shortened down to a minimal size range of 25 to 33-bp, representing ∼30% of extant *P. tetraurelia* IESs. A phylogenetic analysis of IESs across *P. aurelia* species showed that shortening has been accompanied by a switch in their excision mechanism. Indeed, the most recently inserted IESs (*i*.*e*. the youngest) were shown to depend on scnRNAs and heterochromatin marks, while old IESs have become independent of these epigenetic factors (Sellis et al. 2021). Here we show that late excised IESs tend to be the youngest and that, similar to their TE ancestors, their elimination depends on scnRNAs and histone marks. In addition, excision of late IESs tends to depend on the presence of iesRNAs, which have been proposed to boost excision through a positive-feedback loop (Sandoval et al. 2014). The stimulatory contribution of iesRNAs might explain why excision of late IESs precedes imprecise elimination of TEs and other MIC-limited sequences during MAC development. We also report that early excised IESs tend to be the oldest and are enriched for smaller sizes (54.5% belong to the 25 to 33-bp peak). They are also mostly independent of ncRNAs and heterochromatin marks and tend to be the least error-prone. Our data therefore provide experimental support to the proposed evolutionary scenario of *Paramecium* IESs, showing that excision timing reflects their evolutionary age (Fig. 5B). We further provide evidence that IESs have evolved through optimization for efficient excision, combining an early and accurate excision process. Closer analysis of the intrinsic properties of very early excised IESs furthermore revealed a strong nucleotide sequence signal at their ends, which varies according to IES size (TATAG for 25 to 33-bp IESs, TACAG for 42 to 140-bp IESs). In contrast, late excised IESs only exhibit a conserved TA dinucleotide at their ends. These observations suggest that acquisition of a stronger sequence motif has allowed “optimized” IESs to loosen their requirement for sRNAs for excision. Sequence-dependent determination of efficient excision would explain the previous observations that 25 to 33-bp MAC genome segments flanked by terminal TATAG inverted repeats are strikingly under-represented in the somatic MAC genome (Swart et al. 2014), possibly because such sequences are highly excision-prone. MAC genome segments of any size flanked by terminal TACAG inverted repeats are overal poorly represented in the *Paramecium* genome, thus precluding their harmful excision (Swart et al. 2014). The present study therefore points to the joint contribution of IES nucleotide sequence and size as intrinsic determinants for efficient IES excision. Several hypotheses might explain how these intrinsic determinants could work. By facilitating the formation of particular DNA structures, they could help to target specific sequences for elimination, either through a passive mechanism involving nucleosome exclusion to increase their accessibility, or by actively promoting the assembly of the Pgm-endonuclease complex. The different consensus sequences according to IES size might also be related to a distinct spatial organization of IES ends within the excision complex formed for very short *vs* longer IESs (Arnaiz et al. 2012). Another non-exclusive hypothesis might be that conserved sequence motifs locally help to position the Pgm catalytic domain on its cleavage sites.

Studying the *Paramecium* model, with its nuclear dimorphism and ability to precisely excise TE-related IESs even when inserted inside coding sequences, provides a unique opportunity to monitor how TEs have evolved within their host genomes. Further work on *Paramecium* PGR will make it possible to decipher the evolutionary and mechanistic switch from sRNA- and heterochromatin-mediated TEs silencing to efficient physical elimination of TE-related sequences from the genome. The characterization of a set of efficiently excised IESs, which has become independent of sRNAs and the heterochromatin pathway, paves the way to future biochemical studies that will address the longstanding question of how domesticated PiggyBac transposases are recruited to specific DNA cleavage sites to carry out precise DNA excision.

## Methods

### Cell growth and autogamy time-courses

Culture of *P. tetraurelia* wild type 51 new (Gratias and Bétermier 2003) or its mutant derivative 51 *nd7-1* (Dubois et al. 2017) was performed as previously described (Abello et al. 2020). For autogamy time-courses, cells (∼20-30 vegetative fissions) were seeded at a final concentration of 250 cells/mL in inoculated WGP medium with an OD_600nm_ adjusted to 0.1. Autogamy was usually triggered by starvation on the next day. We performed 6 independent autogamy time-courses (tc1 to tc6). For each time-course, The T0 time-point was defined as the time (in hours) when 50% of the cells in the population have a fragmented MAC. We further defined 5 developmental stages: DEV1 (T2.5-T3), DEV2 (T7-T12), DEV3 (T20-T24), DEV4 (T30), DEV5 (T48), with time-points following T0 based on (Arnaiz et al. 2017). At each selected time-point, 0.7-2 L of culture (at a concentration of 1500-3500 cells/mL) were processed for nuclear preparation or 30 mL for whole cell immunofluorescence. To inhibit DNA replication during autogamy, aphidicolin (Sigma-Aldrich, ref. A0781), a specific inhibitor of DNA polymerase α (Ikegami et al. 1978), was added at T2.5 at a final concentration of 15 μM, and the same volume was added a second time at T10. Cells were harvested and nuclei were isolated at T20. For all time-courses, the survival of post-autogamous progeny was tested as described before (Dubois et al. 2017).

### Preparation of nuclei

Nuclear preparations enriched in developing MACs were obtained as previously described (Arnaiz et al. 2012) with few modifications. The cell pellet was resuspended in 6-10 volumes of lysis buffer supplemented with 2x Protease Inhibitor Cocktail Set I (PICS, Cabiochem, ref. 539131), kept on ice for 15 min and disrupted with a Potter-Elvehjem homogenizer (100 to 400 strokes). Lysis efficiency was monitored with a Zeiss Lumar.V12 fluorescence stereo-microscope, following addition of 66 µg/mL DAPI. Nuclei were collected through centrifugation at 1000 g for 2 min and washed four times with 10 volumes of washing buffer. The nuclear pellet was either diluted 2-fold in washing buffer containing glycerol (13% final concentration) and frozen as aliquots at -80°C or diluted 2-fold in washing buffer supplemented with 2x PICS and loaded on top of a 3 mL sucrose (2.1 M) solution layer before ultra-centrifugation in a swinging rotor for 1 h at 210,000 g. After gentle washes, the pellet was resuspended in 1 volume of washing buffer containing glycerol (13% final concentration) and frozen as aliquots at - 80°C.

### Cellular and nuclear immunolabeling

A peptide corresponding to PgmL1 amino acid sequence 1 to 266 and carrying a C-terminal His tag was used for guinea pig immunization (Proteogenix). Sera were purified by antigen affinity purification to obtain highly specific α−PgmL1-GP antibodies (0.8 mg/mL). RNAi targeting the *PGML1* gene during autogamy, immunofluorescence labeling of whole cells and quantification of PgmL1 signal were performed as described previously (Bischerour et al. 2018). Cells were extracted with ice-cold PHEM (60 mM PIPES, 25 mM HEPES, 10 mM EGTA, 2 mM MgCl_2_ pH 6.9) + 1% Triton prior to fixation and immunostaining with α-PgmL1-GP (1:2000). New MAC labeling was adapted from a described method (Sardo et al. 2017). Nuclear preparations were immunostained on ice for 1 h in TBS (10 mM Tris pH 7.4, 0.15 M NaCl) + 3% BSA containing α-PgmL1-GP (1:1000). Nuclei were washed twice in TBS + 3% BSA and stained for 45 min with Alexa Fluor (AF) 488-conjugated goat anti-guinea pig IgG (1:500, ThermoFisher Scientific). Nuclei were finally washed twice in TBS + 3% BSA and resuspended in PI/RNase staining buffer (BD Pharmingen, ref. 550825). All centrifugation steps were performed at 500 g for 1 min at 4°C. Samples were kept in the dark at 4°C until processing.

### Flow cytometry

Stained nuclei were filtered through sterile 30 µm cell strainers (Sysmex filters, CellTrics® ref. 04-004-2326) and processed for flow cytometry. Immunostained nuclei were analyzed on a CytoFlex S cytometer (Beckman Coulter) with a 488 nm laser for scatter measurements (Forward scatter, or FSC, and Side scatter, or SSC) and AF488 excitation, and a 561 nm laser for PI excitation. AF488 and PI staining signals were respectively collected using a 525/40 nm band pass filter and a 610/20 nm band pass filter. Immunostained nuclei were sorted on a Moflow Astrios EQ cell sorter (Beckman Coulter) with a 488-nm laser for scatter measurements (Forward Scatter, or FCS, and Side Scatter, or SSC) and AF488 excitation, and a 561 nm laser for PI excitation. AF488 and PI staining signals were respectively collected using a 526/52 nm band pass filter and a 614/20 nm band pass filter. Phosphate Buffered Saline-like (Puraflow Sheath Fluid, Beckman Coulter) was used as sheath and run at a constant pressure of 10 or 25 PSI. Frequency of drop formation was 26 or 43 kHz. Purify mode was used for sorting in order to reach a maximum rate of purity (>95%). The instrument used a 100 µm nozzle. A threshold on the PI signal was optimized to increase collecting speed (∼1000 events per second). Data were collected using Summit software (Beckman Coulter). Nuclei were first gated based on their Side Scatter (SSC) and high PI signal, and sorted according to their AF488 signal. AF488-positive events were backgated onto SSC *vs* PI to optimize the gating. Doublets were discarded using PI-area and PI-Height signals. Nuclei (<30,000) were collected into 100 μl of Buffer AL (QIAamp DNA Micro Kit, QIAGEN) and immediately lysed by pulse-vortexing. Final volume was adjusted to 200 μL with PI/RNase staining buffer. We confirmed that the FANS procedure yields pure anlagen in an experiment (tc3) in which old MAC fragments contained a marker transgene absent from the anlagen (*Supplemental* Fig. S3; Supplemental Methods).

### Estimation of new MAC DNA content by flow cytometry

Estimation of the absolute DNA content (C-value) in the new MAC populations was based on a previously described method (Bourge et al. 2018). The DNA content was calculated using the linear relationship between the fluorescent signal from the new MAC peaks and a known internal standard (tomato nuclei, *Solanum lycopersicum L*. cv. Montfavet 63-5, 2C=1946 Mbp). Briefly, leaves were chopped with a razor blade in a Petri dish with PI/RNase staining buffer, filtered through 30-µm cell strainers and added in a constant ratio to an aliquot of stained and filtered *Paramecium* nuclei. The C-value (C_new MACs_) for each new MAC subpopulation was calculated using its PI Mean Fluorescence Intensity (MFI_new MACs_), the PI Mean Fluorescence Intensity of the 2C tomato standard (MFI_standard_), and the 2C-value of the tomato standard (2C_standard_):

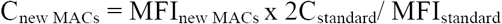

The degree of endoploidy for each new MAC population (C-level) was further estimated by dividing the C-value for each new MAC population by the DNA content of the unrearranged *P. tetraurelia* MIC genome (1C≈100 Mb) (Guérin et al. 2017; Sellis et al. 2021): C-level= C_new MACs_ /1C_mic_

### Genomic DNA extraction and high throughput sequencing

DNA extraction was performed using the QIAamp DNA Micro Kit (Qiagen) as recommended by the manufacturer, with minor modifications. Following a 10-min incubation with proteinase K (2 mg/mL) in buffer AL, the nuclear lysate was directly loaded onto the purification column. Elution was performed with 20-50 μL Buffer AE in DNA LoBind Eppendorf Tubes. DNA concentration was determined using the QBit High Sensitivity kit (Invitrogen) before storage at -20°C. Sequencing libraries were prepared using 1.5 to 8.5 ng of DNA with the TruSeq NGS Library Prep kit from Westburg (WB9024) following manufacturer’s instructions. Alternatively, genomic DNA was fragmented with the S220 Focused-ultrasonicator (Covaris). Fragments were processed with NEBNext Ultra II End Prep Reagents (NEB #E7546) and TruSeq adapters were ligated using the NEBNext Quick Ligation kit (NEB #E6056). Libraries were amplified by PCR using Kapa HiFi DNA polymerase (10-14 cycles). Library quality was checked with an Agilent bioanalyzer instrument (Agilent High Sensitivity DNA kit). Sequencing was performed on 75-75bp paired-end runs, with an Illumina NextSeq500/550 instrument, using the NextSeq 500/550 MID output cycle kit. Demultiplexing was performed with bcl2fastq2-2.18.12 and adapters were removed with Cutadapt 1.15; only reads longer than 10 bp were retained.

### Software and datasets

Standard software, reference genomes and datasets are described in Supplemental Methods. The IES sequence end logos were generated using weblogo (v3.6.0 --composition 0.28 --units bits).

### IES classification

Mapping of sequencing reads on the MAC and the MAC+IES references was used to calculate an IES Excision Score (ES =IES^−^/(IES^+^ + IES^−^) using ParTIES (MIRET default parameters). An ES of 0 means no excision and an ES of 1 means complete IES excision. The violin plots show the distribution of the mean ES score for the two IES boundaries of all IESs. Excision profile classification was carried out on the 44928 annotated IESs, after removing 543 IESs with ES < 0.8, in at least one FRAG sample, which indicates imperfect excision in the old MAC (group defined as " None "). K-means clustering (iter.max=100) was used to define 4 groups based on the ES in all conditions.

### TE and genome coverage

The mean sequencing depth (samtools depth -q 30 -Q 30), normalized by the number of reads mapped on the MIC reference genome, was calculated on TE copies (500 nt min length and localized on MIC contigs > 2kb) and IESs. Only fully mapped reads overlapping at least 4 nucleotides of the annotated feature were considered. As previously described (Guérin et al. 2017), the same window coverage approach was used to estimate genome coverage at each time point. The coverage (multicov -q 30) was calculated for non-overlapping 1-kb windows, then normalized by the total number of mapped reads (RPM). An empirical cutoff of 2.5 RPM was used to decide if the window is covered or not.

### Detection of *de novo* telomere addition sites

*De novo* telomere addition sites were identified on the MIC genome, with the requirement of at least 3 consecutive repeats of either G_4_T_2_ or G_3_T_3_ on mapped reads. A telomere addition site was identified if the read alignment stops at the exact position where the telomeric repeat starts. The number of telomere addition sites was normalized by the number of reads mapped on the MIC genome.

### IES-IES junctions

The ParTIES Concatemer module, developed for this study, was used with default parameters to identify concatemers of excised IESs. Reads were recursively mapped to the IES sequences, as shown in Supplemental Fig. S5A. At each round, reads are mapped to IES sequences and selected if the alignment begins or ends at an IES extremity. If the read is partially aligned, then the unmapped part of the read is re-injected into the mapping and the selection procedure continues until the entire read has been mapped.

### IES excision errors

The ParTIES MILORD module was used with the MAC+IES reference genome to identify IES excision errors. Only error types described in Supplemental Fig. S7A were considered. The number of non-redundant errors was normalized by the number of mapped reads. PCR duplicates were removed using samtools rmdup.

## Data access

The sequencing data generated for this study have been submitted to the ENA database (https://www.ebi.ac.uk/ena/browser/home) under accession number PRJEB49315.

The statistical data, R scripts and raw images have been deposited at Zenodo (https://doi.org/10.5281/zenodo.6534540).

The cytometry data generated in this study have been submitted to the FlowRepository database (http://flowrepository.org/id/RvFrl4FUJTnaAIDEsEqK3MzxKwQZpkfp7yqzGGSco3tuuLfuAHKrPI2fP65KehpH).

## Acknowledgments and funding sources

We would like to thank Cindy Mathon and Pascaline Tirand for their technical assistance and all members of the Bétermier laboratory for stimulating and fruitful discussions. Special thanks to Joël Acker, Julien Bischerour and Linda Sperling for critical reading of the manuscript. This study was supported by intramural funding from the Centre National de la Recherche (CNRS) and by grants from the *Agence Nationale de la Recherche* (ANR-18-CE12-0005-02 & ANR-21-CE12-0019-01) and the *Fondation pour la Recherche Médicale* (FRM EQU202103012766). The present work has benefited from the expertise of the Imagerie-Gif core facility, supported by the *Agence Nationale de la Recherche* (ANR-11-EQPX-0029/Morphoscope, ANR-10-INBS-04/FranceBioImaging, ANR-11-IDEX-0003-02/ Saclay Plant Sciences). We acknowledge the sequencing and bioinformatics expertise of the I2BC High-throughput sequencing facility, supported by *France Génomique* (funded by the French National Program “Investissement d’Avenir” ANR-10-INBS-09).

## Author contributions

C.Z., M.Bourge, M.Bétermier, O.A, V.R.: designed research, analyzed data.

C.Z, K.G., L.E., M.Bourge, N.M., O.A, V.R., Y.J.: performed research.

O.A.: developped bioinformatic pipelines and conducted bioinformatic data analysis.

C.Z., K.G., M.Bourge, M.Bétermier, O.A.,V.R.: wrote the paper.

## References

Abello A, Régnier V, Arnaiz O, Le Bars R, Betermier M, Bischerour J. 2020. Functional diversification of Paramecium Ku80 paralogs safeguards genome integrity during precise programmed DNA elimination. PLoS Genet 16: e1008723.

Allen SE, Hug I, Pabian S, Rzeszutek I, Hoehener C, Nowacki M. 2017. Circular concatemers of ultra-short DNA segments produce regulatory RNAs. Cell 168: 990–999.

Allen SE, Nowacki M. 2020. Roles of Noncoding RNAs in Ciliate Genome Architecture. J Mol Biol. http://www.ncbi.nlm.nih.gov/pubmed/31926952.

Arnaiz O, Mathy N, Baudry C, Malinsky S, Aury JM, Denby Wilkes C, Garnier O, Labadie K, Lauderdale BE, Le Mouel A, et al. 2012. The Paramecium germline genome provides a niche for intragenic parasitic DNA: evolutionary dynamics of internal eliminated sequences. PLoS Genet 8: e1002984.

Arnaiz O, Van Dijk E, Bétermier M, Lhuillier-Akakpo M, de Vanssay A, Duharcourt S, Sallet E, Gouzy J, Sperling L. 2017. Improved methods and resources for paramecium genomics: transcription units, gene annotation and gene expression. BMC Genomics 18: 483.

Aury JM, Jaillon O, Duret L, Noel B, Jubin C, Porcel BM, Segurens B, Daubin V, Anthouard V, Aiach N, et al. 2006. Global trends of whole-genome duplications revealed by the ciliate Paramecium tetraurelia. Nature 444: 171–8.

Baroin A, Prat A, Caron F. 1987. Telomeric site position heterogeneity in macronuclear DNA of Paramecium primaurelia. Nucleic Acids Res 15: 1717–28.

Baudry C, Malinsky S, Restituito M, Kapusta A, Rosa S, Meyer E, Bétermier M. 2009. PiggyMac, a domesticated piggyBac transposase involved in programmed genome rearrangements in the ciliate Paramecium tetraurelia. Genes Dev 23: 2478–2483.

Bennetzen JL, Park M. 2018. Distinguishing friends, foes, and freeloaders in giant genomes. Curr Opin Genet Dev 49: 49–55.

Berger JD. 1973. Nuclear differentiation and nucleic acid synthesis in well-fed exconjugants of Paramecium aurelia. Chromosoma 42: 247–268.

Bétermier M, Bertrand P, Lopez BS. 2014. Is non-homologous end-joining really an inherently error-prone process? PLoS Genet 10: e1004086.

Bétermier M, Borde V, Villartay J-P de. 2020. Coupling DNA Damage and Repair: an Essential Safeguard during Programmed DNA Double-Strand Breaks? Trends in Cell Biology 30: 87– 96.

Bétermier M, Duharcourt S. 2014. Programmed rearrangement in ciliates: Paramecium. Microbiol Spectr 2: MDNA3-0035-2014.

Bétermier M, Duharcourt S, Seitz H, Meyer E. 2000. Timing of Developmentally Programmed Excision and Circularization of Paramecium Internal Eliminated Sequences. Mol Cell Biol 20: 1553– 1561.

Bischerour J, Bhullar S, Denby Wilkes C, Regnier V, Mathy N, Dubois E, Singh A, Swart E, Arnaiz O, Sperling L, et al. 2018. Six domesticated PiggyBac transposases together carry out programmed DNA elimination in Paramecium. Elife 7: e37927.

Bleykasten-Grosshans C, Neuveglise C. 2011. Transposable elements in yeasts. C R Biol 334: 679–86.

Bourge M, Brown SC, Siljak-Yakovlev S. 2018. Flow cytometry as tool in plant sciences, with emphasis on genome size and ploidy level assessment. GenApp 2: 1.

Brennecke J, Aravin AA, Stark A, Dus M, Kellis M, Sachidanandam R, Hannon GJ. 2007. Discrete small RNA-generating loci as master regulators of transposon activity in Drosophila. Cell 128: 1089–103.

Capy P. 2021. Taming, Domestication and Exaptation: Trajectories of Transposable Elements in Genomes. Cells 10. http://www.ncbi.nlm.nih.gov/pubmed/34944100.

Chalker DL, Meyer E, Mochizuki K. 2013. Epigenetics of ciliates. Cold Spring Harb Perspect Biol 5: a017764.

Cheng CY, Orias E, Leu JY, Turkewitz AP. 2020. The evolution of germ-soma nuclear differentiation in eukaryotic unicells. Curr Biol 30: R502–R510.

Choi JY, Lee YCG. 2020. Double-edged sword: The evolutionary consequences of the epigenetic silencing of transposable elements. PLoS Genet 16: e1008872.

Cosby RL, Chang NC, Feschotte C. 2019. Host-transposon interactions: conflict, cooperation, and cooption. Genes Dev 33: 1098–1116.

de Vanssay A, Touzeau A, Arnaiz O, Frapporti A, Phipps J, Duharcourt S. 2020. The Paramecium histone chaperone Spt16-1 is required for Pgm endonuclease function in programmed genome rearrangements. PLoS Genet 16: e1008949.

Déléris A, Berger F, Duharcourt S. 2021. Role of Polycomb in the control of transposable elements. Trends Genet 37: 882–889.

Deniz O, Frost JM, Branco MR. 2019. Regulation of transposable elements by DNA modifications. Nat Rev Genet 20: 417–431.

Dubois E, Bischerour J, Marmignon A, Mathy N, Regnier V, Betermier M. 2012. Transposon invasion of the Paramecium germline genome countered by a domesticated PiggyBac transposase and the NHEJ pathway. Int J Evol Biol 2012: 436196.

Dubois E, Mathy N, Regnier V, Bischerour J, Baudry C, Trouslard R, Betermier M. 2017. Multimerization properties of PiggyMac, a domesticated piggyBac transposase involved in programmed genome rearrangements. Nucleic Acids Res 45: 3204–3216.

Duharcourt S, Keller A-M, Meyer E. 1998. Homology-Dependent Maternal Inhibition of Developmental Excision of Internal Eliminated Sequences in Paramecium tetraurelia. Mol Cell Biol 18: 7075–7085.

Duret L, Cohen J, Jubin C, Dessen P, Goût J-F, Mousset S, Aury J-M, Jaillon O, Noël B, Arnaiz O, et al. 2008. Analysis of sequence variability in the macronuclear DNA of Paramecium tetraurelia: a somatic view of the germline. Genome Res 18: 585–596.

Frapporti A, Miro Pina C, Arnaiz O, Holoch D, Kawaguchi T, Humbert A, Eleftheriou E, Lombard B, Loew D, Sperling L, et al. 2019. The Polycomb protein Ezl1 mediates H3K9 and H3K27 methylation to repress transposable elements in Paramecium. Nat Commun 10: 2710.

Furrer DI, Swart EC, Kraft MF, Sandoval PY, Nowacki M. 2017. Two Sets of Piwi Proteins Are Involved in Distinct sRNA Pathways Leading to Elimination of Germline-Specific DNA. Cell Rep 20: 505–520.

Görtz HD. 1988. Paramecium. Springer-Verlag, Berlin Heidelberg New York.

Gratias A, Bétermier M. 2001. Developmentally programmed excision of internal DNA sequences in Paramecium aurelia. Biochimie 83: 1009–1022.

Gratias A, Bétermier M. 2003. Processing of double-strand breaks is involved in the precise excision of Paramecium IESs. Mol Cell Biol 23: 7152–7162.

Gratias A, Lepere G, Garnier O, Rosa S, Duharcourt S, Malinsky S, Meyer E, Betermier M. 2008. Developmentally programmed DNA splicing in Paramecium reveals short-distance crosstalk between DNA cleavage sites. Nucleic Acids Res 36: 3244–3251.

Guérin F, Arnaiz O, Boggetto N, Denby Wilkes C, Meyer E, Sperling L, Duharcourt S. 2017. Flow cytometry sorting of nuclei enables the first global characterization of Paramecium germline DNA and transposable elements. BMC Genomics 18: 327.

Hamilton EP, Kapusta A, Huvos PE, Bidwell SL, Zafar N, Tang H, Hadjithomas M, Krishnakumar V, Badger JH, Caler EV, et al. 2016. Structure of the germline genome of Tetrahymena thermophila and relationship to the massively rearranged somatic genome. Elife 5. http://www.ncbi.nlm.nih.gov/pubmed/27892853.

Ignarski M, Singh A, Swart EC, Arambasic M, Sandoval PY, Nowacki M. 2014. Paramecium tetraurelia chromatin assembly factor-1-like protein PtCAF-1 is involved in RNA-mediated control of DNA elimination. Nucleic Acids Res 42: 11952–11964.

Ikegami S, Taguchi T, Ohashi M, Oguro M, Nagano H, Mano Y. 1978. Aphidicolin prevents mitotic cell division by interfering with the activity of DNA polymerase-alpha. Nature 275: 458–460.

Kapusta A, Matsuda A, Marmignon A, Ku M, Silve A, Meyer E, Forney JD, Malinsky S, Bétermier M. 2011. Highly Precise and Developmentally Programmed Genome Assembly in Paramecium Requires Ligase IV–Dependent End Joining. PLoS Genet 7. https://www.ncbi.nlm.nih.gov/pmc/articles/PMC3077386/ (Accessed April 10, 2020).

Kapusta A, Suh A, Feschotte C. 2017. Dynamics of genome size evolution in birds and mammals. Proc Natl Acad Sci U S A 114: E1460–E1469.

Ketting RF, Haverkamp TH, van Luenen HG, Plasterk RH. 1999. Mut-7 of C. elegans, required for transposon silencing and RNA interference, is a homolog of Werner syndrome helicase and RNaseD. Cell 99: 133–41.

Klobutcher LA, Herrick G. 1997. Developmental genome reorganization in ciliated protozoa: the transposon link. Progr Nucleic Acid Res Mol Biol 56: 1–62.

Le Mouël A, Butler A, Caron F, Meyer E. 2003. Developmentally regulated chromosome fragmentation linked to imprecise elimination of repeated sequences in paramecia. Eukaryot Cell 2: 1076– 1090.

Lepère G, Bétermier M, Meyer E, Duharcourt S. 2008. Maternal noncoding transcripts antagonize the targeting of DNA elimination by scanRNAs in Paramecium tetraurelia. Genes Dev 22: 1501–1512.

Lepere G, Nowacki M, Serrano V, Gout JF, Guglielmi G, Duharcourt S, Meyer E. 2009. Silencing-associated and meiosis-specific small RNA pathways in Paramecium tetraurelia. Nucleic Acids Res 37: 903–15.

Lhuillier-Akakpo M, Frapporti A, Denby Wilkes C, Matelot M, Vervoort M, Sperling L, Duharcourt S. 2014. Local effect of enhancer of zeste-like reveals cooperation of epigenetic and cis-acting determinants for zygotic genome rearrangements. PLoS Genet 10: e1004665.

Lilly MA, Duronio RJ. 2005. New insights into cell cycle control from the Drosophila endocycle. Oncogene 24: 2765–2775.

Maliszewska-Olejniczak K, Gruchota J, Gromadka R, Denby Wilkes C, Arnaiz O, Mathy N, Duharcourt S, Betermier M, Nowak JK. 2015. TFIIS-Dependent Non-coding Transcription Regulates Developmental Genome Rearrangements. PLoS Genet 11: e1005383.

Marmignon A, Bischerour J, Silve A, Fojcik C, Dubois E, Arnaiz O, Kapusta A, Malinsky S, Betermier M. 2014. Ku-mediated coupling of DNA cleavage and repair during programmed genome rearrangements in the ciliate Paramecium tetraurelia. PLoS Genet 10: e1004552.

Miró-Pina C, Charmant O, Kawaguchi T, Holoch D, Michaud A, Cohen I, Humbert A, Jaszczyszyn Y, Chevreux G, Del Maestro L, et al. 2022. Paramecium Polycomb repressive complex 2 physically interacts with the small RNA-binding PIWI protein to repress transposable elements. Developmental Cell. https://www.sciencedirect.com/science/article/pii/S1534580722002088 (Accessed April 22, 2022).

Preer JR. 1976. Quantitative predictions of random segregation models of the ciliate macronucleus. Genet Res 27: 227–238.

Prescott DM. 1994. The DNA of ciliated protozoa. Microbiol Rev 58: 233–267.

Sandoval PY, Swart EC, Arambasic M, Nowacki M. 2014. Functional diversification of Dicer-like proteins and small RNAs required for genome sculpting. Dev Cell 28: 174–88.

Sardo L, Lin A, Khakhina S, Beckman L, Ricon L, Elbezanti W, Jaison T, Vishwasrao H, Shroff H, Janetopoulos C, et al. 2017. Real-time visualization of chromatin modification in isolated nuclei. J Cell Sci 130: 2926–2940.

Sellis D, Guerin F, Arnaiz O, Pett W, Lerat E, Boggetto N, Krenek S, Berendonk T, Couloux A, Aury JM, et al. 2021. Massive colonization of protein-coding exons by selfish genetic elements in Paramecium germline genomes. PLoS Biol 19: e3001309.

Singh A, Maurer-Alcalá XX, Solberg T, Gisler S, Ignarski M, Swart EC, Nowacki M. 2022. RNA-mediated nucleosome depletion is required for elimination of transposon-derived DNA. 2022.01.04.474918. https://www.biorxiv.org/content/10.1101/2022.01.04.474918v1 (Accessed May 3, 2022).

Singh DP, Saudemont B, Guglielmi G, Arnaiz O, Goût J-F, Prajer M, Potekhin A, Przybòs E, Aubusson-Fleury A, Bhullar S, et al. 2014. Genome-defence small RNAs exapted for epigenetic mating-type inheritance. Nature 509: 447–452.

Swart EC, Denby Wilkes C, Sandoval PY, Hoehener C, Singh A, furrer DI, Arambasic M, ignarski M, Nowacki M. 2017. Identification and analysis of functional associations among natural eukaryotic genome editing components [version 1; peer review: 1 approved, 1 approved with reservations]. F1000Research 6.

Swart EC, Wilkes CD, Sandoval PY, Arambasic M, Sperling L, Nowacki M. 2014. Genome-wide analysis of genetic and epigenetic control of programmed DNA deletion. Nucleic Acids Res 42: 8970–8983.

Tabara H, Sarkissian M, Kelly WG, Fleenor J, Grishok A, Timmons L, Fire A, Mello CC. 1999. The rde-1 gene, RNA interference, and transposon silencing in C. elegans. Cell 99: 123–32.

Zilberman D, Cao X, Jacobsen SE. 2003. ARGONAUTE4 control of locus-specific siRNA accumulation and DNA and histone methylation. Science 299: 716–9.

