## Supplementary Information for "Developmental timing of programmed DNA elimination in *Paramecium tetraurelia* recapitulates germline transposon evolutionary dynamics"

**This PDF file includes:**

| | |
|---|---|
| Supplemental Methods | p2 |
| Supplemental Figures S1 to S12 | p5 |
| Supplemental Tables S1, S2, S6 | p18 |
| Supplemental References | p21 |

### Supplemental methods

#### Transformation of the 51 *nd7-1* strain

A complementing plasmid carrying a functional *ND7* gene was microinjected into the MAC of vegetative 51 *nd7-1* cells to allow for easy screening of transformant clones (Dubois et al. 2017). The complementing *ND7* transgene (Tg *ND7*) bears a silent Single Nucleotide Polymorphism (SNP) (T->C scaffold 51_5: 596316) compared to the endogenous gene (designated as endo *ND7*). Transgene injection levels (Copy Per Haploid Genome or CPHG) were determined by qPCR on genomic DNA extracted from vegetative transformants, using a LightCycler 480 and the LightCycler 480 SYBR green Master Kit (Roche Diagnostics). Couples of oligonucleotide primers (5'TTCCAAGGACCCTGCATAAT3'/5'GGACTGAACCAGAGCCTTCA3') and *(*5'AGCTCCTGATGGCAAGACAT3'/5'AGTCTGTCCACAACCAGCTG3') were used to amplify the *ND7* transgene and genomic *KU80c* reference locus, respectively.

#### Contribution of contaminating old MAC fragments to sorted new MAC DNA

We used FANS to sort nuclei from the Tg *ND7* clone at the DEV2 stage (Supplemental Fig. S3B) and deep-sequenced the DNA extracted from three sorted populations (new MACs, peak 1; new MACs, peak 2; fragments) (tc3 in Supplemental Tables S1, S2**).** For each sample, we experimentally determined the ratio Tg *ND7*/endo *ND7*. For old MAC fragments, this ratio is equal to the transgene CPHG of the parental microinjected clone ($CPHG_{FRAG}$). For the new MAC samples, because sorted new MAC DNA contains DNA from contaminating old MACs, this ratio represents the apparent transgene CPHG ($CPHG_{app}$). For each sample, we counted the copies of the *ND7* locus from Tg *ND7* and/or endo *ND7* and defined the % of contamination as follows:

% Contamination = (endo $ND7_{FRAG}$/ endo $ND7_{sample}$) x100

with:

endo $ND7_{FRAG}$ = Tg *ND7*/ $CPHG_{FRAG}$

endo $ND7_{sampl\ e}$= Tg *ND7*/ $CPHG_{app}$

which gives:

% Contamination = $CPHG_{app}/CPHG_{FRAG}$

We used three independent methods to calculate the transgene CPHG values (Supplemental Fig. S3C). In the first method (qPCR), CPHG values were calculated from a qPCR performed on sequencing libraries using the *ND7* transgene and *KU80c* oligonucleotide primers. In the second method, the SNP present in the *ND7* transgene was used to quantify the number of sequencing reads coming from the endogenous *ND7* locus or from the Tg *ND7* transgene. The ratio between the number of Tg and endo *ND7* reads gives the transgene CPHG in each sample. The third method used genomic read coverage calculated with bamCoverage (v3.2.1 --binsize 20 -smothlength 50 --region scaffold51_5:592000:598000). The ratio between the mean coverage of the *ND7* gene (594873-596396) and the mean coverage of its upstream region (592000-594000), minus 1 gives the transgene CPHG in each sample. Contamination levels within the range of 0.9-6.1% and 1.1-4.7% were obtained for the first and second DNA peaks, respectively, indicative of the high purity of the sorted anlagen.

**Software and R packages**

Sequencing reads were mapped on genome references using Bowtie2 (v2.2.9 --local --X 500). The resulting alignments were analyzed using samtools (v1.9), ParTIES (v1.05 https://github.com/oarnaiz/ParTIES) (Denby Wilkes et al. 2016) and bedtools (v2.26). R (v4) packages were used to generate images (ggplot2 v3.3.5; ComplexHeatmap v2.6.2; GenomicRanges v1.42).

**Reference genomes and datasets**

The paired-end sequencing data were mapped on *P. tetraurelia* strain 51 MAC (ptetraurelia_mac_51.fa), MAC+IES (ptetraurelia_mac_51_with_ies.fa) or MIC (ptetraurelia_mic2.fa) reference genomes (Arnaiz et al. 2012; Guérin et al. 2017). Gene annotation v2.0 (ptetraurelia_mac_51_annotation_v2.0.gff3), IES annotation v1 (internal_eliminated_sequence_PGM_ParTIES.pt_51.gff3) and TE annotation v1.0 (ptetraurelia_mic2_TE_annotation_v1.0.gff3) were used in this study (Arnaiz et al. 2012; Guérin et al. 2017). All files are available from the ParameciumDB download section (https://paramecium.i2bc.paris-saclay.fr/download/Paramecium/tetraurelia/51/) (Arnaiz et al. 2020). DNA sequencing data of *Paramecium* cells depleted of Ezl1, TFIIS4, Dcl2/3 or Dcl5 were previously published (Lhuillier-Akakpo et al. 2014; Maliszewska-Olejniczak et al. 2015; Sandoval et al. 2014). ParTIES (MIRET module) was used to determine the IESs that were significantly retained compared to the control (Denby Wilkes et al. 2016). An IES is considered to be dependent on the depleted factor for its excision (31,505; 20,524; 3,439 and 2,475 IESs sensitive to *EZL1*, *TFIIS4*, *DCL2/3* and *DCL5* RNAi, respectively), if at least one IES boundary in at least one replicate shows significant retention.

**Annotation of IESs in MIC-limited regions**

The MIC-limited genome consists of IESs as well as regions that are imprecisely eliminated. We looked for additional IESs inside each compartment. The ParTIES MILORD module was used to identify frequent non-canonical excision events. Events inside an annotated IES were designated « Internal IESs » and events in imprecisely eliminated MIC-limited regions were designated « Imp IESs ». « Internal IESs » were annotated using the MAC+IES reference genome (scaffolds > 30kb). Only « internal » and « partial internal » deletion events bounded by two TAs were considered (see Supplemental Fig. S7A). A series of filters (minimum size 20 nt; no coverage of the event in FRAG samples; events must be revealed by at least 10 reads in at least 2 time course samples; only the event supported by the maximum number of reads is kept in the case of overlapping events) were applied to provide a list of 226 putative « Internal IESs ». The 167 « Imp IESs » were annotated using the MIC reference genome (contigs > 1kb) and reads that do not map on the MAC or the MAC+IES reference genomes, to select deletion events (length < 5kb) localized in imprecisely eliminated MIC-limited regions. The same filters used for « Internal IES » identification were applied.

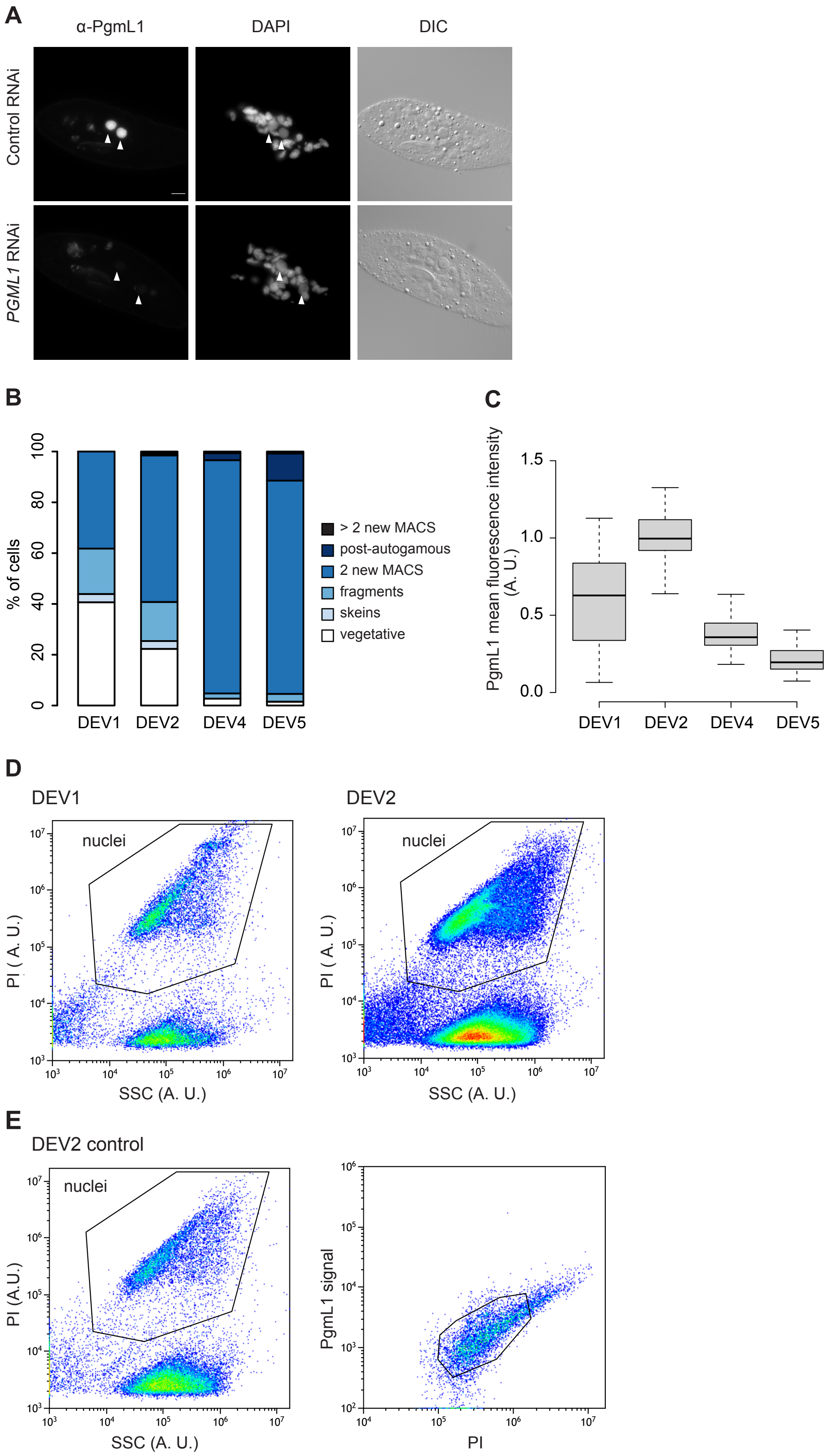

**Supplemental Fig. S1. PgmL1 immunostaining on whole cells and isolated nuclei during developmental time-courses**. (*A*) Validation of the specificity of the α-PgmL1 antibody. Immunostaining of PgmL1 in DEV2 autogamous cells subjected to control or *PGML1* RNAi. (*B*) Cytological progression of autogamy time-course tc1. For each developmental stage (DEV1: T3; DEV2: T10; DEV4: T30; DEV5: T48; with time in hours following T0), the percentage of cells with a defined cytological status (see legend) was determined following DAPI staining of fixed cells and fluorescence microscopy. (C) Boxplot representation of the distribution of PgmL1 fluorescence intensities at the different stages of tc1. 78 to 88 labeled new MACs were quantified at each stage. (D) Flow cytometry analysis of labeled nuclei in tc2. Gating of total nuclei from DEV1 and DEV2 based on their PI and Side Scatter (SSC) signals. (E) Same experiment as in *D* at DEV2 stage without adding the α-PgmL1 primary antibody. A. U.: arbitrary units.

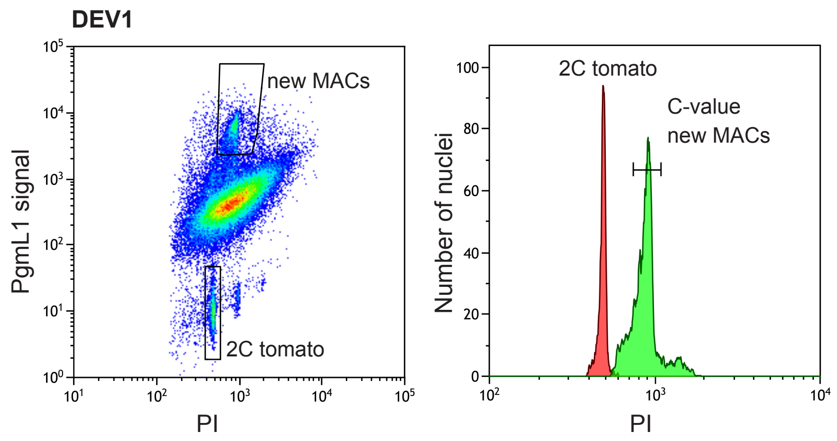

**Supplemental Fig. S2. Example of C-value estimation for sorted new MAC populations.** Tomato nuclei were added to *Paramecium* nuclei isolated at the DEV1 stage from tc4 as shown on the PgmL1 vs PI plot. Peaks of DNA content are defined on the PI histogram. The observed mean PI fluorescence for 2C tomato nuclei is used as an internal standard to calculate the C-value for the new MAC population (see Methods).

**A**

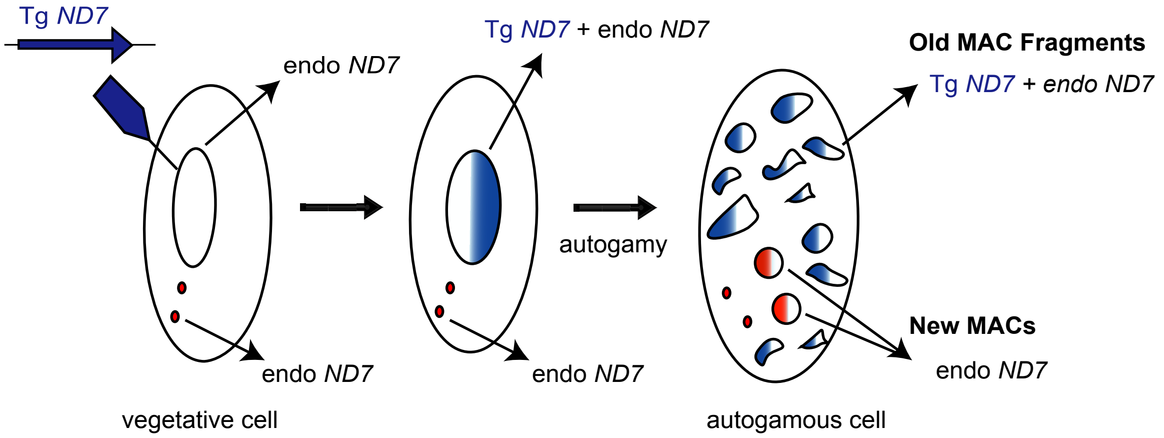

**B**

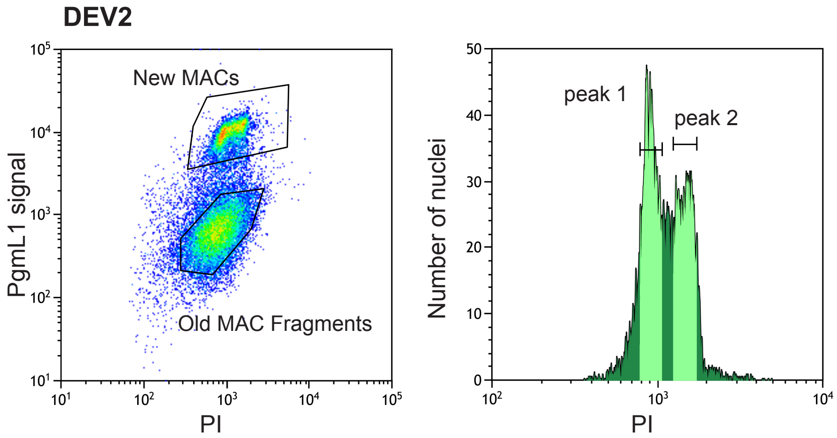

**C**

| | | Fragments | New MACs, peak 1 | New MACs, peak 2 |
|---|---|---|---|---|
| qPCR | CPHG | 183.8 | 1.6 | 2 |
| | **contamination (%)** | | **0.9** | **1.1** |
| ND7 SNP | Reads endo *ND7* | 103 | 115 | 124 |
| | Reads Tg *ND7* | 6198 | 420 | 352 |
| | CPHG | 60.2 | 3.7 | 2.8 |
| | **contamination (%)** | | **6.1** | **4.7** |
| ND7 coverage | *ND7* upstream coverage | 94.9 | 120.5 | 153.9 |
| | *ND7* coverage | 9438 | 654.5 | 604.4 |
| | CPHG | 98.5 | 4.4 | 2.9 |
| | **contamination (%)** | | **4.5** | **3** |

**Supplemental Fig S3. Determination of the contamination level of sorted new MACS with old MAC fragments.** (*A*) Schematic representation of the strategy used. A *Paramecium* clone carrying Tg *ND7* in its somatic MAC was generated by microinjection and starved to induce autogamy. The presence of Tg *ND7* in the DNA isolated from sorted new MACs makes it possible to calculate the contamination by old MAC fragments as described in Supplemental Methods. (*B*) Flow cytometry sorting of nuclei from the Tg *ND7* clone at the DEV2 stage of an autogamy time-course (tc3). Gated events are indicated. Sorted new MAC peaks (1 and 2) are indicated by a light green shading on the PI histogram. Old MAC fragments, sorted as a control, are indicated on the PgmL1 vs PI plots. (*C*) Calculation of the contamination level of purified new MAC DNA from the Tg *ND7* clone. Of note, the qPCR-based method yielded lower contaminating values than the sequencing read counts.

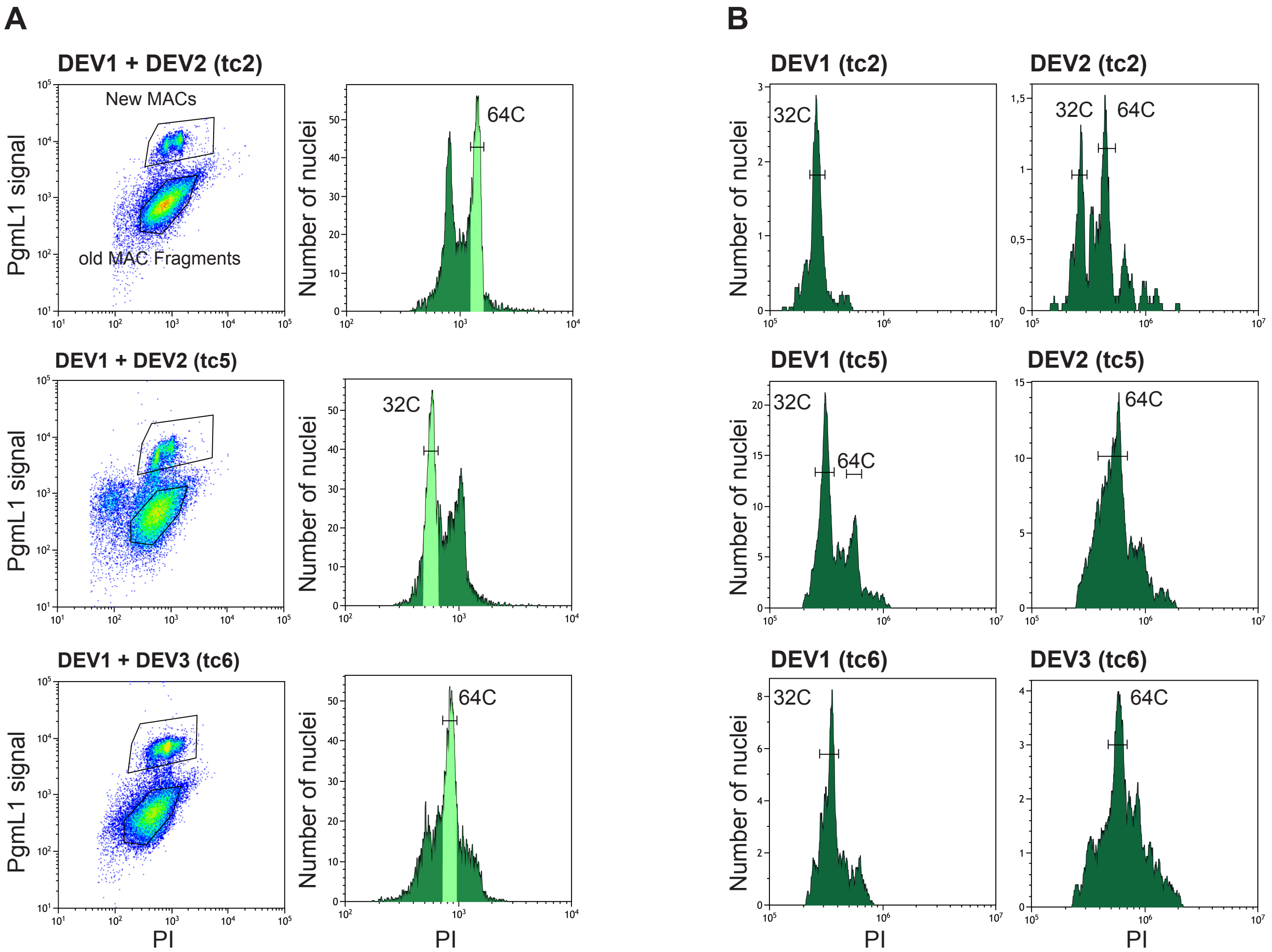

**Supplemental Fig. S4. Plots and histograms for all replicate flow cytometry experiments.** (*A*) Flow cytometry sorting for tc2, tc5 and tc6. The sorting was performed on a pool of nuclei isolated from 2 developmental stages (DEV1 + DEV2 or DEV1 + DEV3). Gated events are indicated. Sorted peaks are indicated by light green shading on the PI histogram. In each case, the sorted new MAC peaks correspond to a unique developmental stage as shown in the analytical flow cytometry presented in *B*. Estimation of the rounds of endoreplication for the indicated peaks is presented in Supplemental Table S1. The sorted old MAC fragments, used as controls, are indicated on the PgmL1 labelling vs PI plots. (*B*) Flow cytometry analysis for tc2, tc5, and tc6. The PI histogram of PgmL1-labelled nuclei is presented for each developmental stage analysed individually. Estimation of the round of endoreplication for each peak was performed using tomato nuclei as an internal standard, as for sorting experiments.

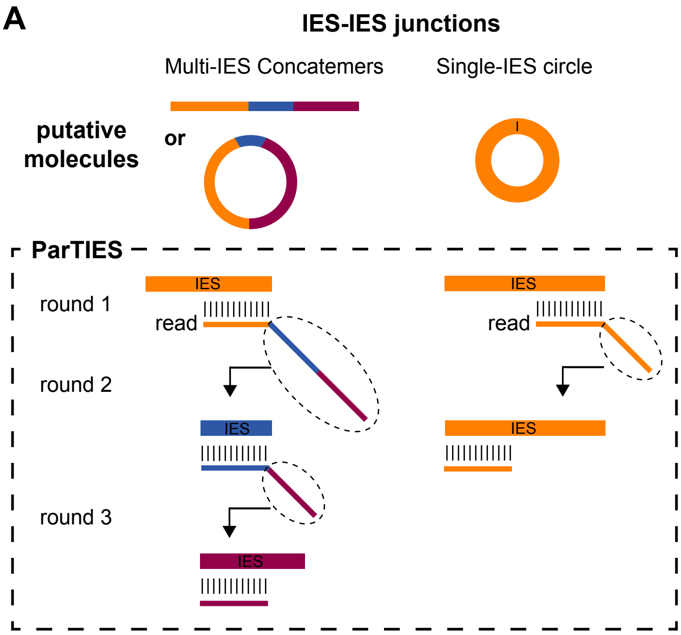

**B**

| | Samples | | Multi-IES concatemers | Single-IES circles | % of single-IES circles |
|---|---|---|---|---|---|
| | FRAG | tc2 | 1.56 | 0 | **0** |
| | | tc3 | 10.73 | 0.06 | **0.6** |
| | | tc4 | 5.46 | 0 | **0** |
| | | tc5 | 4.90 | 0.02 | **0.5** |
| 32C | DEV1 | tc4 | 355.71 | 0.06 | **0** |
| | | tc5 | 708.93 | 0.65 | **0.1** |
| | DEV2 | tc3 | 1127.04 | 4.21 | **0.4** |
| | | tc4 | 1608.79 | 1.18 | **0.1** |
| 64C | | tc3 | 1006.11 | 7.87 | **0.8** |
| | | tc4 | 188.66 | 1.97 | **0.2** |
| | | tc2 | 876.10 | 14.37 | **1.6** |
| | DEV3 | tc6 | 389.98 | 6.68 | **1.7** |
| | | tc4 | 370.58 | 2.62 | **0.7** |
| 128C | DEV3 | tc4 | 54.63 | 1.28 | **2.3** |
| | DEV4 | tc4 | 34.80 | 0.82 | **2.3** |

**C**

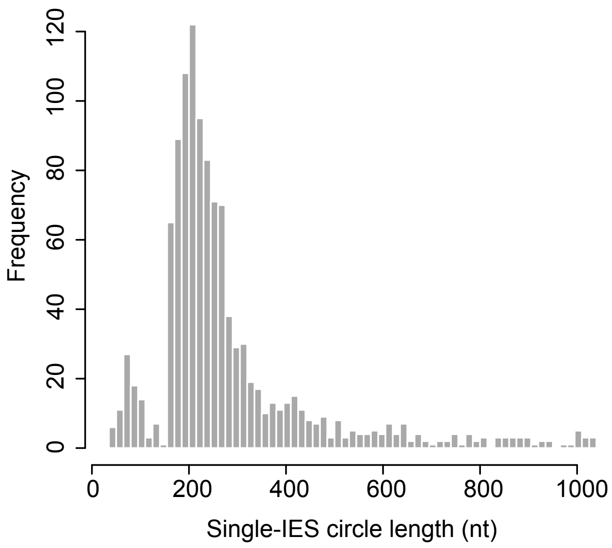

**Supplemental Fig. S5. Detection of IES-IES junctions.** (*A*) Schematic representation of the bioinformatic method used for the detection of IES-IES junctions. The ParTIES concatemer module was developped to detect multi-IES concatemers and single-IES circles, using reads mapped recursively on IES sequences. The procedure is schematically presented in the box and described in Methods. (*B*) Normalized number of sequencing reads (RPM) corresponding to the two types of molecules (multi-IES concatemers or single-IES circles) resulting from the ligation of excised IES ends (see schematic representation in *A*). Linear or circular concatemers cannot be distinguished with Illumina sequencing. (*C*) Size distribution of IESs involved in at least one single-IES circle.

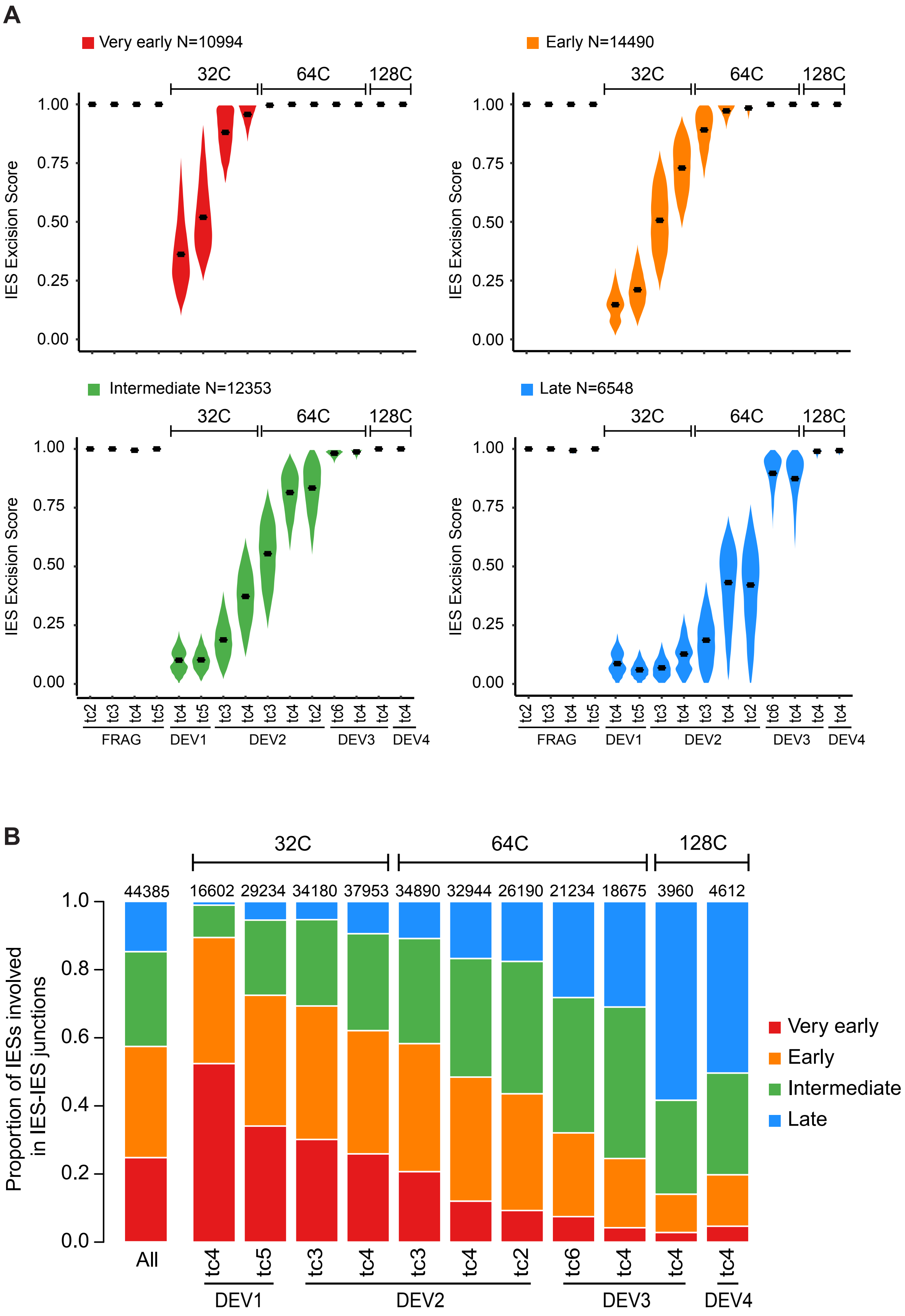

**Supplemental Fig S6. Detection of excision products.** (*A*) ES distribution in each sample for each excision profile group. (*B*) Proportion of IESs involved in at least one excised IES-IES junction in the four excision profile groups (indicated by colors). "All" is the random expectation for all IESs. The number of IESs in each dataset is indicated above each bar.

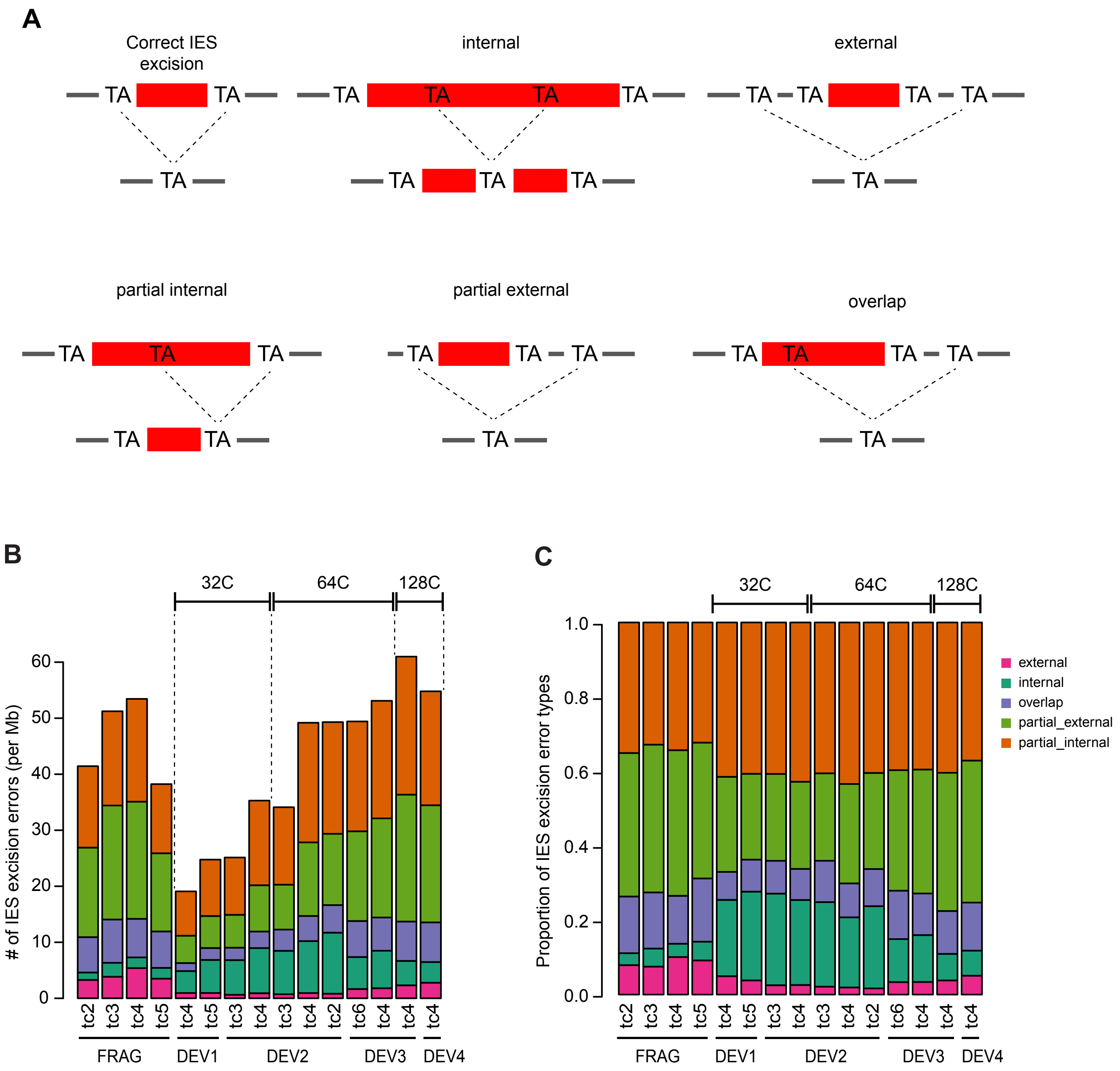

**Supplemental Fig. S7. Detection of IES excision errors.** (*A*) Schematic representation of IES excision errors. In internal or external errors, the two alternative TAs are misplaced: inside or on each flanking side of the reference IES, respectively. Partial internal and partial external excision errors use one correct TA boundary and the other TA inside or outside of the reference IES, respectively. Overlapping error uses one TA inside and the other outside of the reference IES. (*B*) Normalized counts (per Mb mapped on the genome) of excision errors in the different sorted new MAC populations. (*C*) Proportion of the different types of excision errors in the different sorted new MACs or fragments.

**A**

| IES group | Genes (%) | P-value | CDS (%) | P-value | Introns (%) | P-value |
|---|---|---|---|---|---|---|
| All | 84.5 | - | 76 | - | 5.4 | - |
| Very early | 90.3 | 6.7e-55 | 82.6 | 1e-49 | 5.1 | 0.2 |
| Early | 88.4 | 1.4e-31 | 79.4 | 1.4e-17 | 5.8 | 0.1 |
| Intermediate | 83.9 | 0.1 | 74.9 | 0.01 | 5.7 | 0.3 |
| Late | 72.1 | 2.9e-138 | 63.8 | 6.6e-100 | 4.9 | 0.1 |

**B**

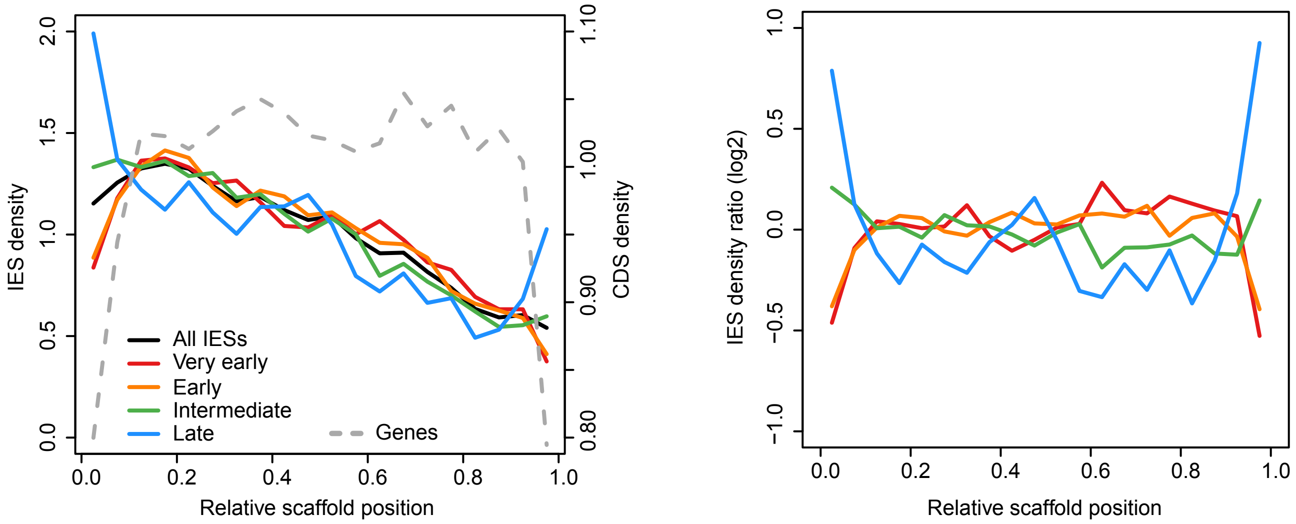

**Supplemental Fig. S8. IES location for excision profile groups.** (*A*) Table showing the percentages of IESs in genes, coding sequences (CDS) and introns for all IESs and for IESs belonging to each group. A chi2 test was performed to compare the proportion obtained in each group with the proportion for all IESs, and the associated p-values are presented. (*B*) IES density along scaffolds. Relative densities for IES site location were calculated (bin of 0.05) on all scaffolds longer than 30 kb, previously oriented to have higher IES density on the first left quarter of the scaffold. Gene density was calculated using the same method and the middle position of each coding exon. Left panel: the gray dotted line shows gene density and the black curve shows IES density. The red, orange, green and light blue curves show the IES density for the 4 groups of excision profiles. Right panel: ratio between IES density for each group relative to the total IES density.

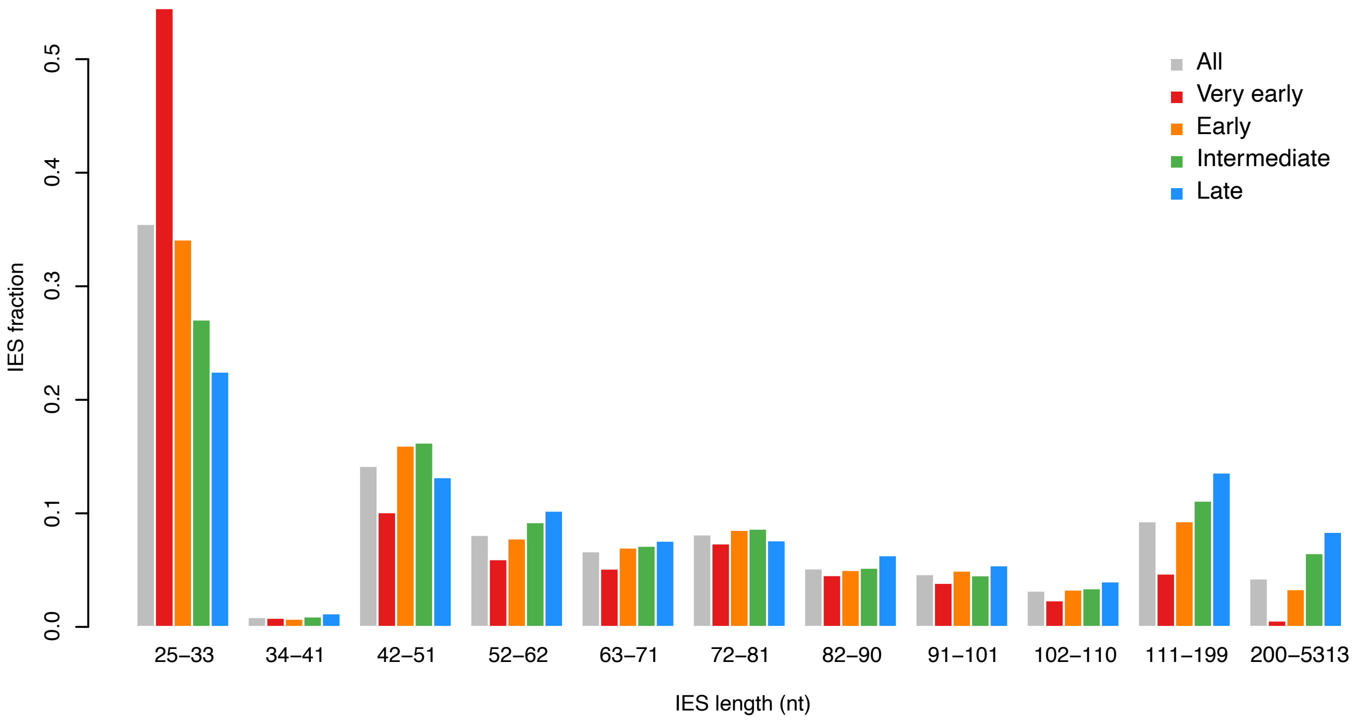

**Supplemental Fig. S9.** IES fraction in each IES length peak, for all IESs (gray) and for IESs belonging to the four excision profile groups.

**A**

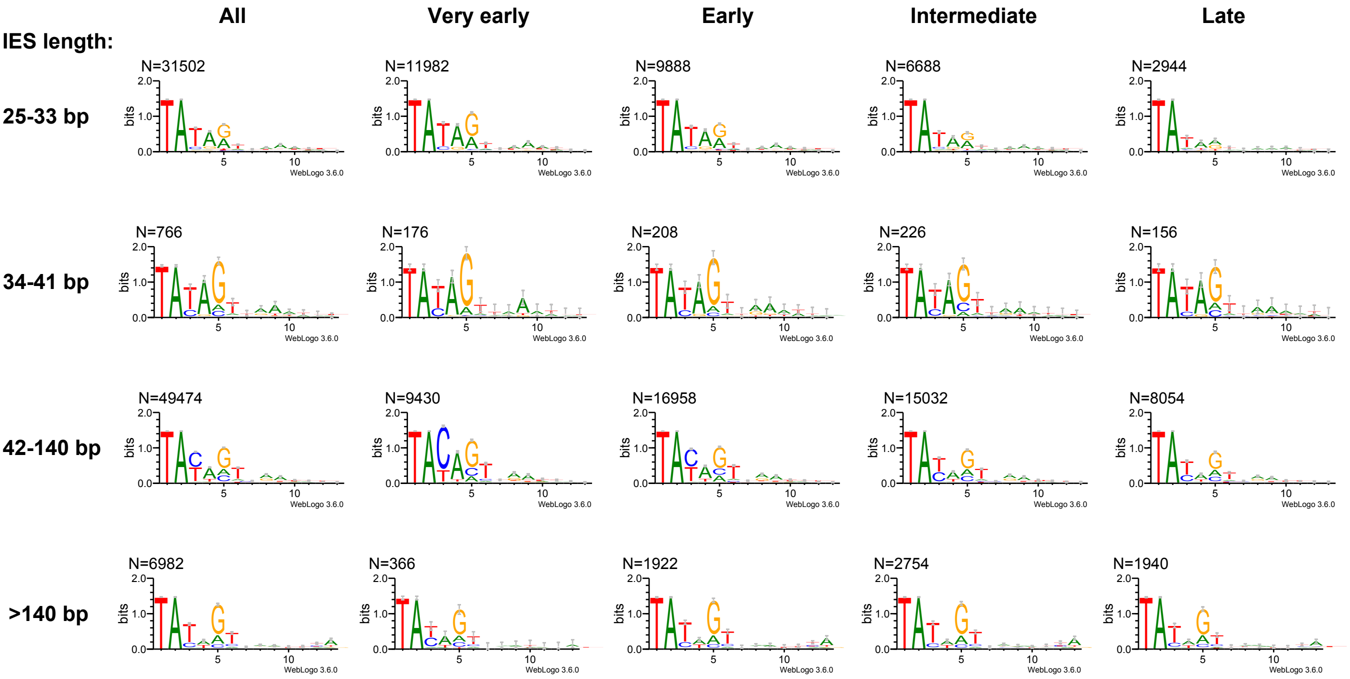

**B**

| | pos. | base | All | Very early | Late |
|---|---|---|---|---|---|
| **25-33 bp** | 3 | T | 79% | 82%** | 72% |
| | 4 | A | 76% | 82%** | 65% |
| | 5 | G | 52% | 62%*** | 35% |
| **42-140 bp** | 3 | C | 49% | 77%**** | 30% |
| | 4 | A | 73% | 88%*** | 62% |
| | 5 | G | 60% | 64% | 57% |

**Supplemental Fig. S10. Analysis of IES ends.** (*A*) Sequence logos of IES ends, calculated according to the IES size (rows) and excision profile group (columns). The first column is for all IESs. (*B*) Base frequencies at positions 3, 4, 5 of IES ends for all, very early and late IESs from the 25-33 bp and 42-150pb length categories. The frequencies for very early and late IESs were compared using a statistical test using a binomial frequency test (binom.test) and a Pearson-Klopper exact method to calculate the upper bound of the confidence interval (binom.confint with the parameters conf.level=0.95). Resulting p-values were adjusted for multiple testing using the Benjamini & Hochberg method. Frequencies with a p-value < 0.001 are considered as significantly different (p-values: * <= 1e-50; ** <1e-100; *** < 1e-300; **** < 1e-400).

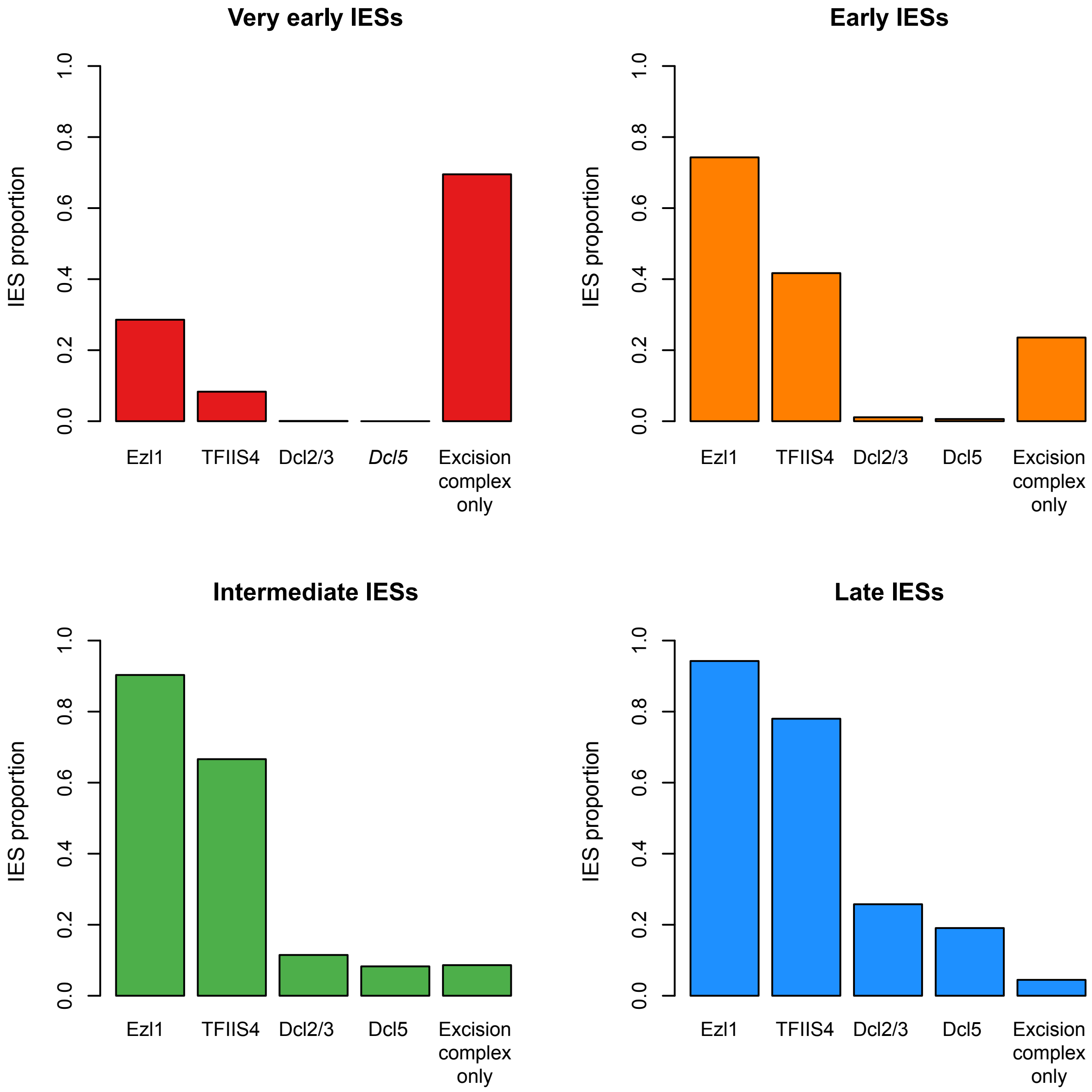

**Fig. S11. Proportion of Ezl1, TFIIS4, Dcl2/3 or Dcl5-dependent IESs compared to the proportion of IESs only dependent upon Pgm (labelled "excision complex only").** Each excision profile group is represented in a separate panel.

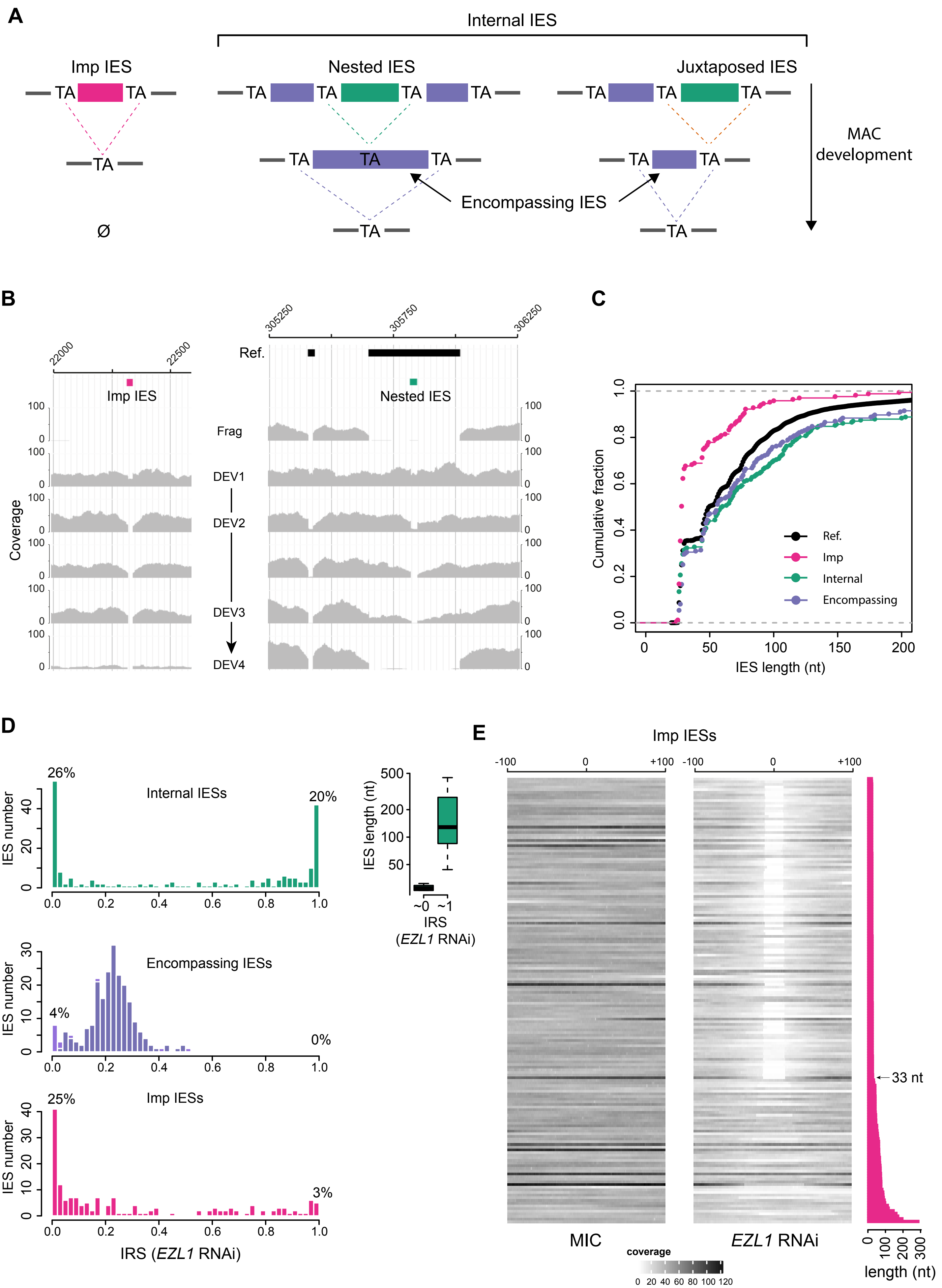

A
Internal IES
Imp IES
Nested IES
Juxtaposed IES
MAC development
Encompassing IES
B
Imp IES
Nested IES
Ref.
Frag
DEV1
DEV2
DEV3
DEV4
Coverage
C
Cumulative fraction
IES length (nt)
Ref.
Imp
Internal
Encompassing
D
Internal IESs
Encompassing IESs
Imp IESs
IES number
IRS (EZL1 RNAi)
IES length (nt)
IRS
(EZL1 RNAi)
E
Imp IESs
MIC
EZL1 RNAi
coverage
33 nt
length (nt)

**Supplemental Fig. S12. Annotation and properties of IESs localized in MIC-specific regions.** (*A*) Schematic representation of excision events transiently observed during MAC development. These newly characterized TA-bound IESs can be of two types: IESs localized in imprecisely eliminated regions that will be completely removed from the mature MAC (Imp IES) or sequences localized within an IES of the reference set (internal IES). In the latter case, two situations can be distinguished depending on whether the new IES is excised between fully internal TAs (nested IES) or shares one TA with its encompassing IES (juxtaposed IES). (*B*) Genome browser screenshots showing one Imp IES (left panel, genome location: NODE_7045_length_27888_cov_20.982180:22000-22500) and one nested IES (right panel, genome location: scaffold51_106_with_IES: 305250-306250) that was already described inside IES 51A6649 from the $A^{51}$ surface antigen gene. The histogram tracks correspond to the sequencing coverage in the sorted samples from the tc4 time-course (DEV1 32C, DEV2 32C, DEV2 64C, DEV3 64C, DEV4 128C). (*C*) Cumulative fraction of the characterized IES datasets according to their length (nt). Ref.: reference set of IESs; Imp: Imp IESs; Internal: Internal IESs (nested or juxtaposed); Encompassing: Encompassing IESs. (*D*) Histogram distributions (bin 0.02) of IES Retention Scores (IRS) in *EZL1* RNAi for IESs belonging to the following categories: Imp (N=167), Internal (N=226), encompassing (N=223). Only IESs with an IRS calculated from at least 10 reads are displayed (N=164, 209, 223 respectively). The percentage of IESs for the two extreme bins (IRS around 0 and IRS around 1) of the distribution are indicated above the bars. For Internal IESs, the length distribution is added as a boxplot for these two IRS categories. (*E*) Heatmaps of coverage within a 200-nt window around Imp IESs (0 being the center of each Imp IES) for a vegetative MIC DNA sample (left) (Sellis et al. 2021) and DNA after *EZL1* RNAi (right) (Lhuillier-Akakpo et al. 2014). IESs, on each row, have been ordered according to their lengths, as illustrated on the pink histogram. The gray scale color intensity corresponds to sequencing depth coverage.

**Supplemental Table S1. Analysis of the peaks of DNA content in all flow cytometry sorting experiments performed in this study.**

| Time course | Stage | PI Fluorescence Peak | Mean fluorescence (± SD) | N | CV | Range of C-value (Mb) | Range of C-level | Estimated C-level |
|---|---|---|---|---|---|---|---|---|
| tc2 | DEV2 | tomato | 458.78 ± 18.21 | 1154 | 3.97 | | | |
| | | new MACs | 1359.06 ± 94.96 | 1509 | 6.99 | 5157.2-6422.4 | 52 - 64 | ~64 |
| tc3 | DEV2 | tomato | 583.9 ± 30.61 | 1922 | 5.24 | | | |
| | | new MACs, peak 1 | 979.07 ± 96.94 | 1762 | 9.9 | 2793.5-3784.5 | 28 - 38 | ~32 |
| | | new MACs, peak 2 | 1585.39 ± 127.26 | 1166 | 8.03 | 4617.5-6023.6 | 46 - 60 | ~64 |
| tc4 | DEV1 | tomato | 484.53 ± 21.31 | 1949 | 4.4 | | | |
| | | new MACs | 878.66 ± 69.82 | 1692 | 7.95 | 3111.7-3984.6 | 31 - 40 | ~32 |
| | DEV2 | tomato | 481.14 ± 20.74 | 1544 | 4.31 | | | |
| | | new MACs, peak 1 | 945.2 ± 49.49 | 995 | 5.24 | 3473-4204.3 | 35 - 42 | ~32 |
| | | new MACs, peak 2 | 1643.69 ± 98.33 | 1302 | 5.98 | 5992-7363.1 | 60 - 74 | ~64 |
| | DEV3 | tomato | 479.92 ± 19.44 | 1037 | 4.05 | | | |
| | | new MACs, peak 1 | 1666.59 ± 103.88 | 629 | 6.2 | 6089.9-7482.1 | 61 - 75 | ~64 |
| | | new MACs, peak 2 | 2801.07 ± 218.55 | 3024 | 7.8 | 10064-12761 | 101 - 128 | ~128 |
| | DEV4 | tomato | 479.34 ± 17.77 | 1134 | 3.71 | | | |
| | | new MACs | 3327.55 ± 623.38 | 4200 | 18.73 | 10585.8-16657.3 | 106-167 | ~128 |
| tc5 | DEV1 | tomato | 358.59 ± 17.7 | 1794 | 4.94 | | | |
| | | new MACs | 615.47 ± 51.65 | 985 | 8.39 | 2915.8-3808.3 | 29 - 38 | ~32 |
| tc6 | DEV3 | tomato | 569.46 ± 30.9 | 4324 | 5.43 | | | |
| | | new MACs | 1612.44 ± 209.23 | 7796 | 12.98 | 4548.3-6582.3 | 46 - 66 | ~64 |
| Aphi | DEV1 | tomato | 500.3 ± 28.59 | 2664 | 5.71 | | | |
| | | new MACs | 824.06 ± 86.57 | 1115 | 9.38 | 2713.5-3756.7 | 27 - 38 | ~32 |
| | DEV3 DMSO | tomato | 489.86 ± 29.25 | 3348 | 5.97 | | | |
| | | new MACs | 1639.53 ± 124.76 | 3223 | 7.61 | 5678.5-7453.8 | 57- 75 | ~64 |
| | DEV3 Aphi | tomato | 482.78 ± 30.2 | 4678 | 6.25 | | | |
| | | new MACs | 878.58 ± 82.28 | 1794 | 9.36 | 3020.8-4131.5 | 30 - 41 | ~32 |

N: number of nuclei counted. CV : coefficient of variation of the DNA peaks (CV%=SD /mean fluorescence x100). The C-value and C-level for each new MAC population are estimated as described in *Methods* and as shown in *Supplemental* Fig. S2. They are presented as an interval based on the standard deviation observed for each peak distribution (tomato standard and new MACs). The last column shows the estimated C-level, corresponding to multiples of rounds of replication.

**Supplemental Table S2.** Sequencing data with ENA accession numbers.

| Sample | ENA Accession | # of sequenced reads | # of reads mapped on the MAC | % of reads mapped on the MAC | # of reads mapped on the MIC | % of reads mapped on the MIC |
|---|---|---|---|---|---|---|
| FRAG (tc2) | ERS9193158 | 93676108 | 90415500 | 97% | 92174955 | 98% |
| FRAG (tc3) | ERS9193159 | 77662786 | 74896931 | 96% | 76186388 | 98% |
| FRAG (tc4) | ERS9193160 | 65870282 | 63439898 | 96% | 64934908 | 99% |
| FRAG (tc5) | ERS9193161 | 91547362 | 87757956 | 96% | 89260601 | 98% |
| 32C-DEV1 (tc4) | ERS9193162 | 66916976 | 54543630 | 82% | 65616314 | 98% |
| 32C-DEV1 (tc5) | ERS9193163 | 78936684 | 60877696 | 77% | 76636518 | 97% |
| 32C-DEV2 (tc3) | ERS9193164 | 122760576 | 95220091 | 78% | 121176641 | 99% |
| 32C-DEV2 (tc4) | ERS9193165 | 76951432 | 61309368 | 80% | 75772371 | 98% |
| 64C-DEV2 (tc3) | ERS9193166 | 148205894 | 119409760 | 81% | 145926265 | 98% |
| 64C-DEV2 (tc4) | ERS9193167 | 74494670 | 61925119 | 83% | 73833316 | 99% |
| 64C-DEV2 (tc2) | ERS9193168 | 102849336 | 84030260 | 82% | 101352683 | 99% |
| 64C-DEV3 (tc6) | ERS9193169 | 92185998 | 77383722 | 84% | 90327042 | 98% |
| 64C-DEV3 (tc4) | ERS9193170 | 80484180 | 69241353 | 86% | 79497333 | 99% |
| 128C-DEV3 (tc4) | ERS9193171 | 54960378 | 50024855 | 91% | 54345156 | 99% |
| 128C-DEV4 (tc4) | ERS9193172 | 97332452 | 90725576 | 93% | 96665100 | 99% |
| FRAG (Aphi) | ERS9193173 | 102454236 | 99678917 | 97% | 101637829 | 99% |
| 32C-DEV1 (Aphi) | ERS9193174 | 61535736 | 50774640 | 83% | 60975841 | 99% |
| DEV3-DMSO | ERS9193175 | 57173842 | 49637234 | 87% | 56678340 | 99% |
| DEV3-Aphi | ERS9193176 | 81337060 | 69504744 | 85% | 80644613 | 99% |

**Supplemental Table S6.** Retention statistics for several IES categories in different RNAi experiments (*PGM*, *EZL1*, *TFIIS4*, *DCL2/3* and *DCL5)*.

| Category | Number | RNAi | Covered | | | % Not covered |
|---|---|---|---|---|---|---|
| | | | % Retained | % Not Retained | % Uncertain | |
| Reference IESs | 44928 | *PGM* | 99.3 | 0.7 | 0 | 0 |
| | | *EZL1* | 70.3 | 29.7 | 0 | 0 |
| | | *TFIIS4* | 45.9 | 54.1 | 0 | 0 |
| | | *DCL23* | 7.7 | 92.3 | 0 | 0 |
| | | *DCL5* | 5.7 | 94.3 | 0 | 0 |
| Internal | 226 | *PGM* | 87.2 | 0 | 12.8 | 0 |
| | | *EZL1* | 37.2 | 35.8 | 19.5 | 7.5 |
| | | *TFIIS4* | 3.5 | 38.9 | 22.1 | 35.4 |
| | | *DCL23* | 2.2 | 12.4 | 7.1 | 78.3 |
| | | *DCL5* | 0 | 22.6 | 2.7 | 74.8 |
| Internal (Nested) | 120 | *PGM* | 87.5 | 0 | 12.5 | 0 |
| | | *EZL1* | 40 | 33.3 | 20.8 | 5.8 |
| | | *TFIIS4* | 3.3 | 37.5 | 25.8 | 33.3 |
| | | *DCL23* | 3.3 | 15.8 | 10 | 70.8 |
| | | *DCL5* | 0 | 25 | 3.3 | 71.7 |
| Internal (Juxtaposed) | 106 | *PGM* | 86.8 | 0 | 13.2 | 0 |
| | | *EZL1* | 34 | 38.7 | 17.9 | 9.4 |
| | | *TFIIS4* | 3.8 | 40.6 | 17.9 | 37.7 |
| | | *DCL23* | 0.9 | 8.5 | 3.8 | 86.8 |
| | | *DCL5* | 0 | 19.8 | 1.9 | 78.3 |
| Encompassing | 223 | *PGM* | 99.6 | 0.4 | 0 | 0 |
| | | *EZL1* | 93.7 | 6.3 | 0 | 0 |
| | | *TFIIS4* | 72.2 | 27.8 | 0 | 0 |
| | | *DCL23* | 13.9 | 86.1 | 0 | 0 |
| | | *DCL5* | 22.4 | 77.6 | 0 | 0 |
| Internal (bigger than encompassing) | 133 | *PGM* | 87.2 | 0 | 12.8 | 0 |
| | | *EZL1* | 55.6 | 14.3 | 22.6 | 7.5 |
| | | *TFIIS4* | 6 | 31.6 | 31.6 | 30.8 |
| | | *DCL23* | 3.8 | 6 | 9.8 | 80.5 |
| | | *DCL5* | 0 | 30.1 | 4.5 | 65.4 |
| Internal (smaller than encompassing) | 93 | *PGM* | 87.1 | 0 | 12.9 | 0 |
| | | *EZL1* | 10.8 | 66.7 | 15.1 | 7.5 |
| | | *TFIIS4* | 0 | 49.5 | 8.6 | 41.9 |
| | | *DCL23* | 0 | 21.5 | 3.2 | 75.3 |
| | | *DCL5* | 0 | 11.8 | 0 | 88.2 |
| Imp | 167 | *PGM* | 97 | 0 | 1.8 | 1.2 |
| | | *EZL1* | 14.4 | 54.5 | 29.3 | 1.8 |
| | | *TFIIS4* | 1.8 | 33.5 | 17.4 | 47.3 |
| | | *DCL23* | 0.6 | 82.6 | 7.2 | 9.6 |
| | | *DCL5* | 0 | 2.4 | 0 | 97.6 |

The first column indicates the IESs under study (reference IESs: all previously annotated IESs) and the second column indicates their numbers. The IES Retention Score (IRS) was used to determine whether an IES is retained. For IES categories “Reference IESs” and “Encompassing”, the previously described statistical test was used (Denby Wilkes et al. 2016). For the other categories, retention was examined only when the IES is covered by at least 10 reads ("covered IES"). Then, IESs with an IRS > 0.8 are qualified as "retained", IRS< 0.2 as "not retained" and between 0.2 and 0.8, as "uncertainly retained". The table shows the percentage of IESs falling into each category.

### Supplemental References

Arnaiz O, Mathy N, Baudry C, Malinsky S, Aury JM, Denby Wilkes C, Garnier O, Labadie K, Lauderdale BE, Le Mouel A, et al. 2012. The Paramecium germline genome provides a niche for intragenic parasitic DNA: evolutionary dynamics of internal eliminated sequences. *PLoS Genet* **8**: e1002984.

Arnaiz O, Meyer E, Sperling L. 2020. ParameciumDB 2019: integrating genomic data across the genus for functional and evolutionary biology. *Nucleic Acids Res* **48**: D599–D605.

Denby Wilkes C, Arnaiz O, Sperling L. 2016. ParTIES: a toolbox for Paramecium interspersed DNA elimination studies. *Bioinformatics* **32**: 599–601.

Dubois E, Mathy N, Regnier V, Bischerour J, Baudry C, Trouslard R, Betermier M. 2017. Multimerization properties of PiggyMac, a domesticated piggyBac transposase involved in programmed genome rearrangements. *Nucleic Acids Res* **45**: 3204–3216.

Guérin F, Arnaiz O, Boggetto N, Denby Wilkes C, Meyer E, Sperling L, Duharcourt S. 2017. Flow cytometry sorting of nuclei enables the first global characterization of Paramecium germline DNA and transposable elements. *BMC Genomics* **18**: 327.

Lhuillier-Akakpo M, Frapporti A, Denby Wilkes C, Matelot M, Vervoort M, Sperling L, Duharcourt S. 2014. Local effect of enhancer of zeste-like reveals cooperation of epigenetic and cis-acting determinants for zygotic genome rearrangements. *PLoS Genet* **10**: e1004665.

Maliszewska-Olejniczak K, Gruchota J, Gromadka R, Denby Wilkes C, Arnaiz O, Mathy N, Duharcourt S, Betermier M, Nowak JK. 2015. TFIIS-Dependent Non-coding Transcription Regulates Developmental Genome Rearrangements. *PLoS Genet* **11**: e1005383.

Sandoval PY, Swart EC, Arambasic M, Nowacki M. 2014. Functional diversification of Dicer-like proteins and small RNAs required for genome sculpting. *Dev Cell* **28**: 174–88.

Sellis D, Guerin F, Arnaiz O, Pett W, Lerat E, Boggetto N, Krenek S, Berendonk T, Couloux A, Aury JM, et al. 2021. Massive colonization of protein-coding exons by selfish genetic elements in Paramecium germline genomes. PLoS Biol 19: e3001309.
